# Evolution tunes functional sub-state interconversion to boost enzyme function

**DOI:** 10.64898/2025.12.31.697001

**Authors:** Daniel Salamonsen, Andrea Pierangelini, Karol Buda, Daojiong Wang, Marc W. van der Kamp, Nobuhiko Tokuriki, H. Adrian Bunzel, Christopher Frøhlich

## Abstract

Enzymes do not operate as static structures, but continuously fluctuate between different conformations. Enzymes therefore dynamically sample conformations with varying catalytic activity. However, it remains largely unexplored whether evolution can exploit the conformational dynamics between sub-states to improve activity. Here, we dissect the evolutionary trajectory of the β-lactamase OXA-48 toward improved ceftazidime hydrolysis. Evolution relieved conformational bottlenecks by promoting alternate functional sub-states, gradually shifting the rate-limiting step from substrate binding to sub-state interconversion, and finally to the chemical step. Reorganization of the conformational landscape enhanced OXA-48’s ability to hydrolyze ceftazidime and introduced a trade-off in its native activity against meropenem. This trade-off stemmed from catalytic incompatibility between the native and the evolved sub-state populations. Our findings highlight the transitions between functional sub-states as a mechanism of natural selection, shaping functional divergence and offering new strategies for enzyme and antibiotic engineering.

## INTRODUCTION

Evolution can yield enzymes that accelerate chemical reactions with remarkable efficiency. While contemporary evolutionary theory often focuses on static structural effects to explain how evolution improves function, proteins are inherently dynamic. Enzymes sample a range of conformational sub-states, each of which may differ in catalytic activity and together determine their overall efficiency.^1–6^ Thus, it has been proposed that evolution not only optimizes enzyme activity through static structural changes, but also by shifting conformational ensembles toward functional sub-states with superior activity.^7–11^ Despite numerous structural studies describing protein conformational ensembles and their changes through evolution, experimental data demonstrating how the evolution of functional sub-states improves enzyme activity remain limited.^10,12^

Conformational changes may restrict enzyme activity when structural transitions along the reaction coordinate are rate-limiting.^1,13^ For instance, enzymes such as kinases and dihydrofolate reductase convert between open and closed states to allow substrate binding and exclude solvent during the chemical step.^14–16^ However, emerging evidence suggests that conformational changes orthogonal to the reaction coordinate can likewise modulate activity by populating catalytically superior functional sub-states.^10,17^ Thus, rate-limiting transitions between functional sub-states, in addition to their equilibrium alone, may restrict activity and offer a previously underappreciated mechanism by which evolution can tune enzyme function.^12,18–20^

We recently subjected the β-lactamase OXA-48 to directed evolution, increasing ceftazidime resistance in *Escherichia coli* by 43-fold through five sequential mutations (F72L→S212A→T213A→A33V→K51E, Q5, Tab. S1 and Fig. S1).^21^ Increased resistance coincided with the emergence of a pronounced super-stoichiometric burst during substrate turnover (Fig. S1). While single-turnover bursts typically report on a fast step followed by a slower one, super-stoichiometric bursts indicate that the enzyme transitions to less active sub-states over multiple turnovers.^20,22^ Such bursts may arise from substrate-induced conformational changes, which are commonly described by two mechanisms: (*i*) “conformational selection”, where the substrate binds to a pre-existing state, thereby shifting the equilibrium toward that conformation; and (*ii*) “induced fit”, where substrate binding to one state triggers a conformational change to an alternative conformation (Fig. 1a).^23^ While the emergence of super-stoichiometric bursts in OXA-48 suggests that evolution shifted the equilibrium of functional sub-states, the mechanism by which this was achieved remained unclear (Fig. 1b). Addressing this gap could uncover how conformational ensembles shape enzymatic function, and how evolution can tune the conformational and reaction coordinates to improve activity.

**Fig. 1:**
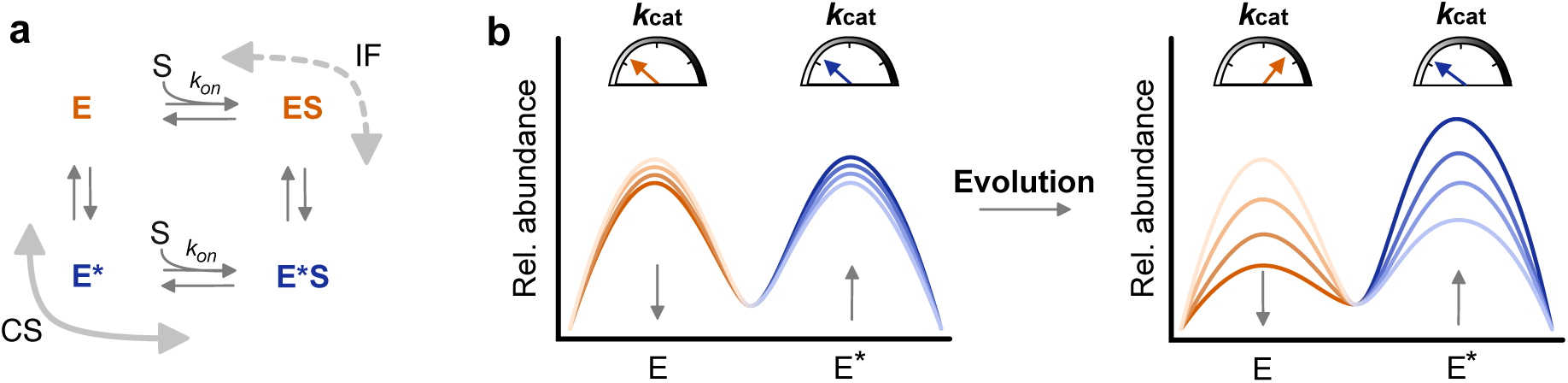
Substrate-induced conformational transitions as drivers of catalytic efficiency. **a.** Induced fit (IF) and conformational selection (CS) are possible mechanisms for substrate (S)-induced conformational changes in enzymes (E). In IF, substrate binding triggers a conformational change from ES to E*S. In CS, the substrate binds preferentially to E*, thereby stabilizing this pre-existing conformation. **b.** Evolution may enhance catalytic efficiency by tuning the activity and population equilibrium of E and E*. One way this can occur is by exploiting substrate-induced shifts in the E ⇌ E* equilibrium (arrows and light-to-dark colors indicate conformational change upon substrate addition). Super-stoichiometric bursts are expected if E is more active than E* and substrate binding induces a conformational change from E to E*.

Here, we demonstrate how evolution affected the energy landscape of OXA-48 through two key mutations (F72L and S212A), which increase OXA-48-mediated ceftazidime resistance by 11-fold (Fig. S1a and Tab. S1).^21^ Through detailed kinetic and structural analyses, we reveal how alleviating kinetic bottlenecks between functional sub-state transitions enhances enzyme activity. Moreover, we demonstrate that shifts in the functional sub-states induce activity trade-offs for an alternative substrate. Our findings reveal that evolution can tune both the interconversion rates between functional sub-states as well as those along the reaction coordinate to optimize enzymatic function. These findings expand our understanding of natural evolution and the emergence of resistance, and provide a framework for exploiting functional sub-states in enzyme engineering and drug design.

## RESULTS

### Evolution of functional sub-state populations

Based on our previously generated evolutionary trajectory of OXA-48,^21^ we interrogated wtOXA-48 and key mutants which conferred an 11-fold increase in ceftazidime resistance: F72L, S212A, and F72L/S212A (Fig. 2a, Fig. S1a, and Tab. S1). To confirm that all variants show super-stoichiometric bursts, we measured ceftazidime turnover kinetics in detail (Tab. 1, Fig. 2b, and Fig. S2a). During evolution from wtOXA-48 to F72L/S212A, the *k*_cat_ of the burst phase (*k*_cat,burst_) increased by 19-fold from 0.008 to 0.15 s^–1^, while the *k*_cat_ of the steady state (*k*_cat,steady_) remained unchanged. In addition, the substrate-dependent inactivation rate (*k*_inact_) increased from >0.004 to 0.04 s^−1^. Importantly, the turnover rates of the burst and steady-state phases converge at decreasing CAZ concentrations, and the burst phase becomes less pronounced (Fig. S2a). This substrate-dependent deactivation should therefore be irrelevant under experimental selection conditions, where substrate concentrations (≤ 14 µM) were much lower than those in the *in vitro* kinetic analysis (≤ 400 µM).^21^ Thus, evolution enhanced activity under physiologically relevant conditions while introducing collateral enzyme inactivation at elevated substrate concentrations. Since the evolution of super-stoichiometric burst kinetics indicate that evolution targeted sub-state interconversions, we next investigated how the catalytic cycle was rewired to access more active sub-states (Fig. 2a).

**Fig. 2:**
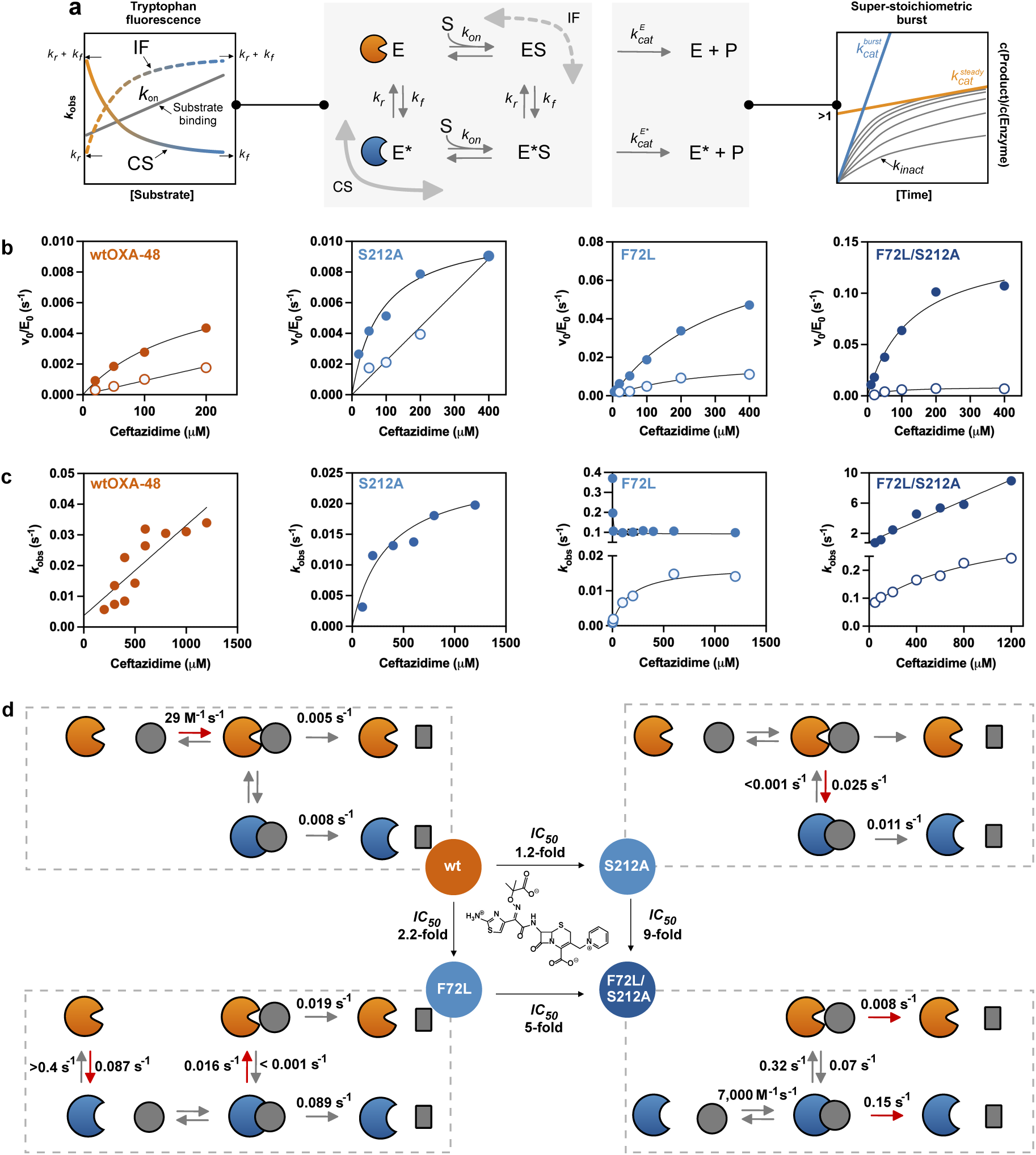
Overview of kinetic measurements and model for ceftazidime hydrolysis. **a.** Substrate binding, induced fit (IF), and conformational selection (CS) mechanisms can affect enzyme activity. Changes in tryptophan fluorescence upon substrate addition (*k*obs) can be monitored to determine pre-steady state kinetics, including substrate binding (*k*on) and transition rates between functional sub-states (*k*f and *k*r). Comparing these pre-steady-state parameters (*k*on, *k*f and *k*r) with substrate turnover (*k*cat,burst and *k*cat,steady) can shed light on the evolution of functional sub-states. **b.** Michaelis-Menten burst (filled circles) and steady-state (empty circles) kinetics (see Fig. S2a for raw traces). **c.** Substrate-binding kinetics were recorded using tryptophan fluorescence (see Fig. S2b for raw traces). Determined *k*obs for the fast (filled circles) and slow (empty circles) phases are shown. **d.** Our kinetic data (Tab. 1) suggest that the enzyme’s main conformation E (orange) is in equilibrium with a secondary conformation E* (blue; substrate: grey sphere; product: grey rectangle). Evolution shifted the equilibrium from E to E*, thereby changing the rate-determining step from substrate binding to the chemical reaction. Rate-determining steps are indicated with red arrows. Ceftazidime *IC50* fold changes refer to the respective changes compared to wtOXA-48 (Tab. S1).

**Tab. 1:** Kinetic parameters for ceftazidime.

|  | wtOXA-48 | S212A | F72L | F72L/S212A |
| --- | --- | --- | --- | --- |
| $k_{\text{cat, burst}} (\text{s}^{-1})$ | $0.008 \pm 0.001$ | $0.011 \pm 0.001$ | $0.089 \pm 0.007$ | $0.15 \pm 0.02$ |
| $k_{\text{cat}}/K_{\text{M, burst}} (\text{M}^{-1} \text{s}^{-1})$ | $4.5 \times 10^1$ | $1.3 \times 10^2$ | $2.6 \times 10^2$ | $1.2 \times 10^3$ |
| $k_{\text{cat, steady}} (\text{s}^{-1})$ | $0.005 \pm 0.001$ | ND | $0.019 \pm 0.002$ | $0.008 \pm 0.001$ |
| $k_{\text{cat}}/K_{\text{M, steady}} (\text{M}^{-1} \text{s}^{-1})$ | $1.0 \times 10^1$ | $2.2 \times 10^1$ | $7.5 \times 10^1$ | $1.4 \times 10^2$ |
| $k_{\text{inact}} (\text{s}^{-1})$ | $>0.004$ | ND | $>0.012$ | 0.04 |
| $k_{\text{on}} (\text{M}^{-1} \text{s}^{-1})$ | $29 \pm 6$ | ND | ND | $7,000 \pm 600$ |
| $k_{\text{r,fast}} (\text{s}^{-1})$ | ND | ND | $>0.4$ | ND |
| $k_{\text{f,fast}} (\text{s}^{-1})$ | ND | ND | $0.087 \pm 0.011$ | ND |
| $k_{\text{r,slow}} (\text{s}^{-1})$ | ND | $<0.001$ | $<0.001$ | $0.07 \pm 0.01$ |
| $k_{\text{f,slow}} (\text{s}^{-1})$ | ND | $0.025 \pm 0.005$ | $0.016 \pm 0.001$ | $0.32 \pm 0.06$ |

ND = Not detectable.
Errors reflect the standard error of the mean based on at least n=2.

To assess how evolution altered conformational dynamics, we mixed enzymes with ceftazidime and monitored responses in tryptophan fluorescence as a readout for conformational changes (Fig. 2c and Fig. S2b). We hypothesized that three distinct binding models can be followed: simple binding (Eq. 1), conformational selection (Eq. 2), or induced fit (Eq. 3). These models can be distinguished from each other based on their binding kinetics, as they display linear, inverse hyperbolic, and hyperbolic relationships between observed rates (*k*_obs_) and [S], respectively (Fig. 2a). For both conformational selection and induced fit, substrate-induced transitions between sub-states are described by their forward and reverse rates (*k*_f_ and *k*_r_), respectively. By comparing binding with turnover kinetics (*k*_cat,burst_ and *k*_cat,steady_), we aimed to examine how evolutionary changes in sub-state interconversion rates are linked to shifts in the rate-limiting step and improved activity.

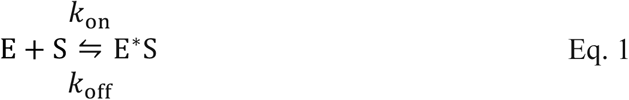

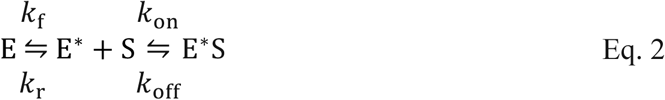

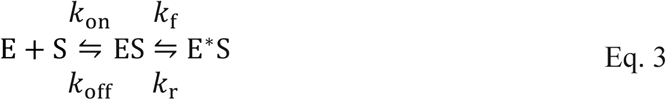

For wtOXA-48, a linear relationship between the *k*_obs_ values for substrate-induced changes in fluorescence and [S] was observed, with a binding rate (*k*_on_) of 29 M^−1^ s^−1^ (Fig. 2c, Tab. 1 and Fig. S2). Thus, turnover is likely limited by substrate binding in wtOXA-48, as *k*_on_ is close to *k*_cat_/*K*_M_ (10 M^−1^ s^−1^). In contrast, the *k*_obs_ values for S212A followed an induced-fit model, saturating at higher ceftazidime concentrations with a *k*_f_ of 0.025 s^−1^. Because *k*_f_ is similar to *k*_cat,burst_ (0.011 s^−1^), a substrate-induced conformational change is likely limiting turnover during the burst phase of S212A (Fig. 2d).

Interestingly, F72L displayed double-exponential binding kinetics (Fig. S2b). The faster phase followed a conformational selection model, indicating that the free enzyme equilibrates between sub-states E and E*, with the substrate binding selectively to E* (Fig. 2d). The forward rate of this interconversion (*k*_f,fast_) matched the *k*_cat_ of the burst phase of F72L (0.087 s^−1^ *versus* 0.089 s^−1^, Tab. 1), suggesting that this conformational change limits the overall turnover during the burst phase. The slower second phase followed an induced fit model with a *k*_f,slow_ = 0.016 s^−1^ and *k*_r,slow_ = 0.001 s^−1^. The forward reaction rate is consistent with *k*_cat,steady_ (0.019 s^−1^), suggesting that the overall rate under steady state conditions is limited by the relaxation from E*S to ES (Fig. 2d). Thus, population of the more productive E*S state likely reflects a branching point within the catalytic cycle, resulting in the super-stoichiometric burst observed in Michaelis-Menten kinetics.

Similar to F72L, the double mutant F72L/S212A displayed biphasic pre-steady-state kinetics (Fig. S2b). However, the fast phase followed a linear relationship between *k*_obs_ and [S], reflecting direct substrate binding (Fig. 2c). The determined *k*_on_ was 6-fold higher than the *k*_cat_/*K*_M_ of the burst phase (7,000 M^−1^ s^−1^ *versus* 1,200 M^−1^ s^−1^, Tab. 1), indicating efficient substrate binding to the more productive E* state (Fig. 2d). The slow phase of F72L/S212A followed an induced fit model with a *k*_f,slow_ = 0.32 s^−1^ and *k*_r,slow_ = 0.07 s^−1^ (Tab. 1). Similar to F72L, the E*S complex interconverts to the less productive ES during turnover, resulting in the observed super-stoichiometric burst phase kinetics.

Overall, directed evolution progressively remodeled the conformational landscape of OXA-48 (Fig. 2d). While catalysis in wtOXA-48 is limited by substrate binding, F72L and S212A unlock and promote more productive functional sub-states. The interconversion between unproductive and productive functional sub-states is likely limiting turnover in these single-point mutants. F72L/S212A overcomes this bottleneck by shifting the equilibrium from E to E*, thereby bypassing this rate-limiting conformational change altogether. Interestingly, the free E* state of F72L/S212A still re-equilibrates to the less productive E state during turnover, which causes the super-stoichiometric burst, and the kinetic bottleneck observed at elevated ceftazidime concentrations.

### Sub-state specialization causes activity trade-offs

To further probe the energy landscape, we studied OXA-48 with an alternative substrate, meropenem, for which binding is unlikely to be rate-limiting.^24^ The fitness landscape comprising all possible combinations of the mutations in Q5 revealed a strong trade-off in *IC_50_* values between ceftazidime and meropenem (Tab. S1 and Fig. S3). This trade-off was primarily driven by F72L, which lowers meropenem resistance by 12-fold while increasing ceftazidime resistance 2-fold (Fig. S3). Given the contrasting trends for ceftazidime and meropenem activity, we reasoned that meropenem could serve as an orthogonal mechanistic probe of the conformational landscape.

To assess ensemble activity for meropenem, we assayed turnover kinetics for wtOXA-48, S212A, F72L, and Q3 (F72L/S212A/T213A). In contrast to ceftazidime, meropenem turnover showed no super-stoichiometric bursts, arguing against substrate-induced perturbations of the conformational ensemble (Fig. 3a and Fig. S2c). F72L, the primary trade-off driver, reduced *k*_cat_ by 14-fold (0.13 s^−1^ to 0.009 s^−1^, Tab. 2), closely matching its 12-fold reduction in meropenem *IC*_50_. Overall, *k*_cat_ values correlated strongly with *IC*_50_ determinations across all tested variants (Pearson correlation, *R*² = 0.95, p = 0.02; Fig. S3). Meropenem resistance is therefore unlikely to be influenced by substrate-induced population shifts and instead determined by the equilibrium distribution of sub-states.

**Fig. 3:**
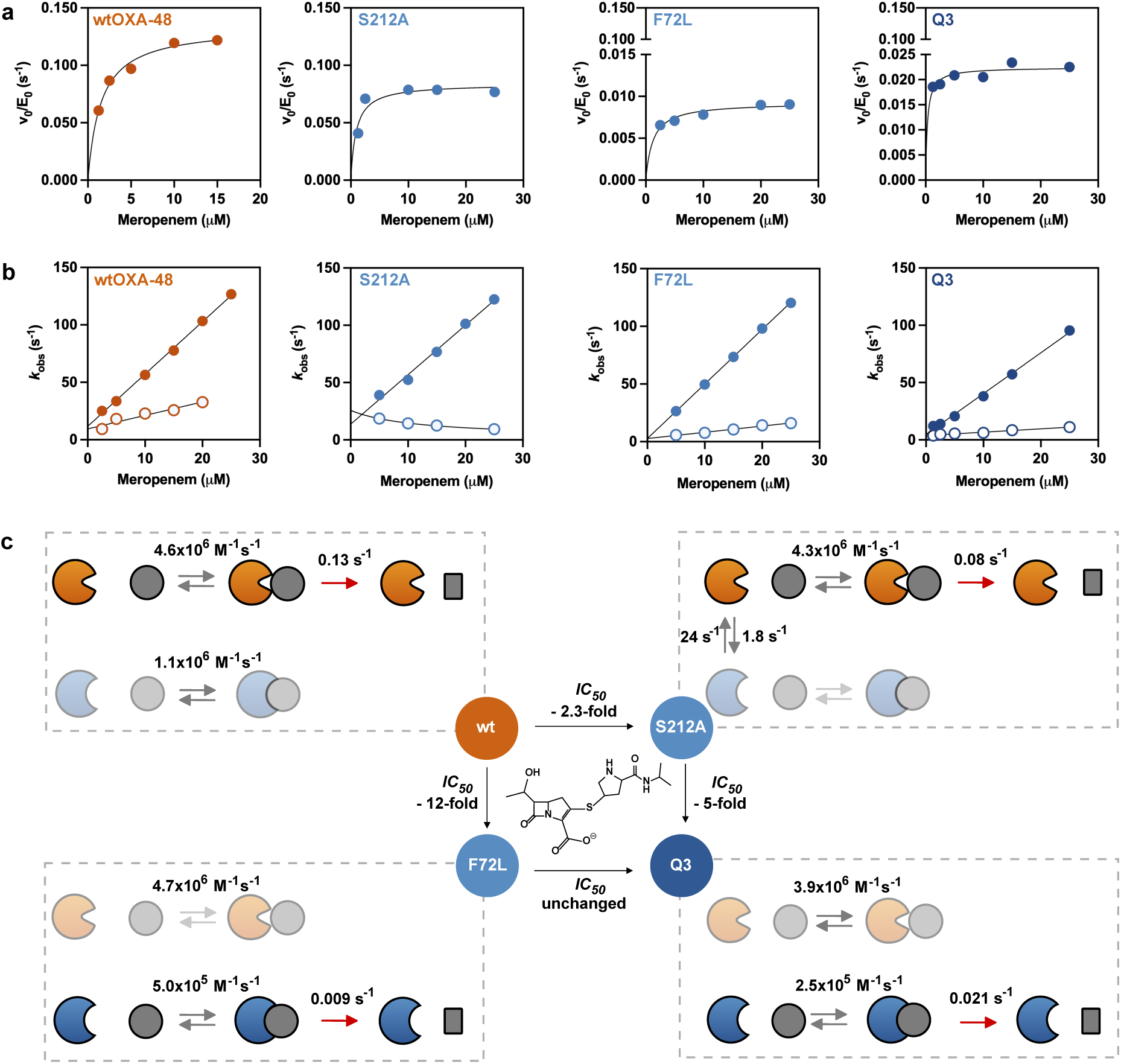
Overview of kinetic measurements and model for meropenem hydrolysis. **a.** Michaelis-Menten kinetics for meropenem (see Fig. S2c for raw traces). **b.** Substrate-binding kinetics were recorded using tryptophan fluorescence (see Fig. S2d for raw traces). Determined *k*obs for the fast (filled circles) and slow (empty circles) phases are displayed. **c.** Proposed reaction mechanism model and rate-determining steps (red arrows) based on the observed kinetic parameters (Fig. S2 and Tab. 2). Fold changes between variants refer to their meropenem *IC50* values (Tab. S1). The enzyme’s main conformation E (orange) is in equilibrium with a secondary conformation E* (blue; substrate: grey sphere; product: grey rectangle). Evolution stepwise shifted the equilibrium from the acylation-efficient E to the acylation-deficient E*. Acylation-deficiency is introduced by F72L and further retained in Q3 by additional mutations. The rate-determining step does not change throughout evolution. Instead, turnover slows down due to a shift in the conformational equilibrium toward E*. Catalytically inferior states are displayed transparently.

**Tab. 2:** Kinetic parameters for meropenem.

|  | wtOXA-48 | S212A | F72L | Q3 |
| --- | --- | --- | --- | --- |
| $k_{\text{cat}} (\text{s}^{-1})$ | $0.13 \pm 0.01$ | $0.08 \pm 0.01$ | $0.009 \pm 0.001$ | $0.021 \pm 0.001$ |
| $k_{\text{cat}}/K_{\text{M}} (\text{M}^{-1} \text{s}^{-1})$ | $8.7 \times 10^4$ | $8.9 \times 10^4$ | $>6.0 \times 10^3$ | $>1.4 \times 10^5$ |
| $k_{\text{on, fast}} (\text{M}^{-1} \text{s}^{-1})$ | $4.6 \times 10^6$ | $4.3 \times 10^6$ | $4.7 \times 10^6$ | $3.9 \times 10^6$ |
| $k_{\text{on, slow}} (\text{M}^{-1} \text{s}^{-1})$ | $1.1 \times 10^6$ | <sup>a</sup> | $5.0 \times 10^5$ | $2.5 \times 10^5$ |
| $k_2 (\text{s}^{-1})$ | 0.07 | 0.08 | Fast: 0.4 (35%)/<br>Slow: 0.009 (65%) | 0.02 |
| Single-turnover<br>conversion | 50% | 50% | 70% | 50% |

<sup>a</sup> The slow phase of S212A followed conformational selection behavior with $k_{\text{r}} = 24 \text{ s}^{-1}$ and $k_{\text{f}} = 1.8 \text{ s}^{-1}$
Errors reflect the standard error of the mean based on at least n=2.

Because meropenem turnover kinetics did not report on the sub-state populations, we next probed the conformational ensemble using time-resolved tryptophan fluorescence (Fig. S2d). All variants exhibited double-exponential fluorescence signals, consistent with two enzyme sub-states. Binding kinetics followed linear models, except for the slow phase of S212A which displayed conformational selection behavior (Fig. 3b). The extracted *k*_on_ values exceeded *k*_cat_/*K*_M_ by 10-100-fold and sub-state interconversion rates of S212A were 23-fold higher than *k*_cat_, confirming that neither meropenem binding nor sub-state interconversion is rate-limiting (Tab. 2). In agreement with the turnover data, binding kinetics thus argue for the presence of two sub-state populations rather than a meropenem-induced perturbation of the conformational ensemble.

To further assess the catalytic competence of the two functional sub-states, we studied enzyme acylation (*k*_2_) with meropenem using single-turnover kinetics (Tab. 2 and Fig. S2e). For wtOXA-48, S212A, and Q3, single-turnover kinetics followed single-exponential curves, although these variants converted only half of the added meropenem. Consistent with the binding data, this indicates the presence of two binding-competent sub-states, only one of which undergoes acylation. In contrast, F72L displayed biphasic acylation, with approximately 70% substrate turnover. The two phases likely stem from two acylating states that may reflect the evolutionary transition between E and E*. E (35%, *k*_2, fast_ = 0.4 s^−1^) and E* (65%, *k*_2, slow_ = 0.009 s^−1^) of F72L exhibited distinct acylation rates, where *k*_2_ of E* limits the overall turnover (*k*_cat_ = 0.009 s^−1^) of the reaction and likely introduces the meropenem trade-off (Tab. 2). The fact that the acylation rate in Q3 (*k*_2_ = 0.02 s^−1^) remained lower than for the wild-type enzyme (*k*_2_ = 0.07 s^−1^), coupled with a uniform acylation population, supports the idea that the combination of F72L and S212A drives a shift in the enzyme equilibrium from E to E* (Fig. 3c and Fig. S2e).

Our kinetic analysis confirms that evolution shifted the conformational equilibrium in OXA-48. The trade-off between ceftazidime and meropenem resistance arises primarily from selection of a ceftazidime-optimized functional sub-state that is poorly suited for carbapenem acylation. Together, these findings highlight the crucial role of the conformational landscape in determining enzyme efficiency and specificity (Fig. 3c).

### Sub-state dynamics shape substrate specificity

To assess whether the functional sub-states inferred from kinetics have a resolvable structural basis, we analyzed crystal structures of apo wtOXA-48 and Q5, the ceftazidime-bound Q5 complex, and meropenem-bound complexes of wtOXA-48 and Q4 (PDB IDs: 4S2P, 8PEB, 8PEC, 6P98, 9T8V, Tab. S2 and Fig. S4).^21,25,26^ Across static crystal structures, no substantial conformational changes were observed for the catalytic serine S70 or the general base K73. Instead, structural data combined with ensemble refinement indicate increased Ω-loop flexibility along the evolutionary trajectory (Fig. 4a). While this loop is typically in a closed conformation in wtOXA-48,^27^ substrate binding in the evolved variants leads to pronounced disorder in the Ω-loop, precluding reliable refinement (Fig. S4). Despite increased flexibility in the Ω-loop, meropenem binding poses in wtOXA-48 and in Q4 remain highly similar. Thus, crystallography captured differences in local flexibility consistent with substrate preferences but, due to the inherently static nature of crystallographic models, did not resolve the conformational dynamics underlying functional sub-state evolution.

**Fig. 4:**
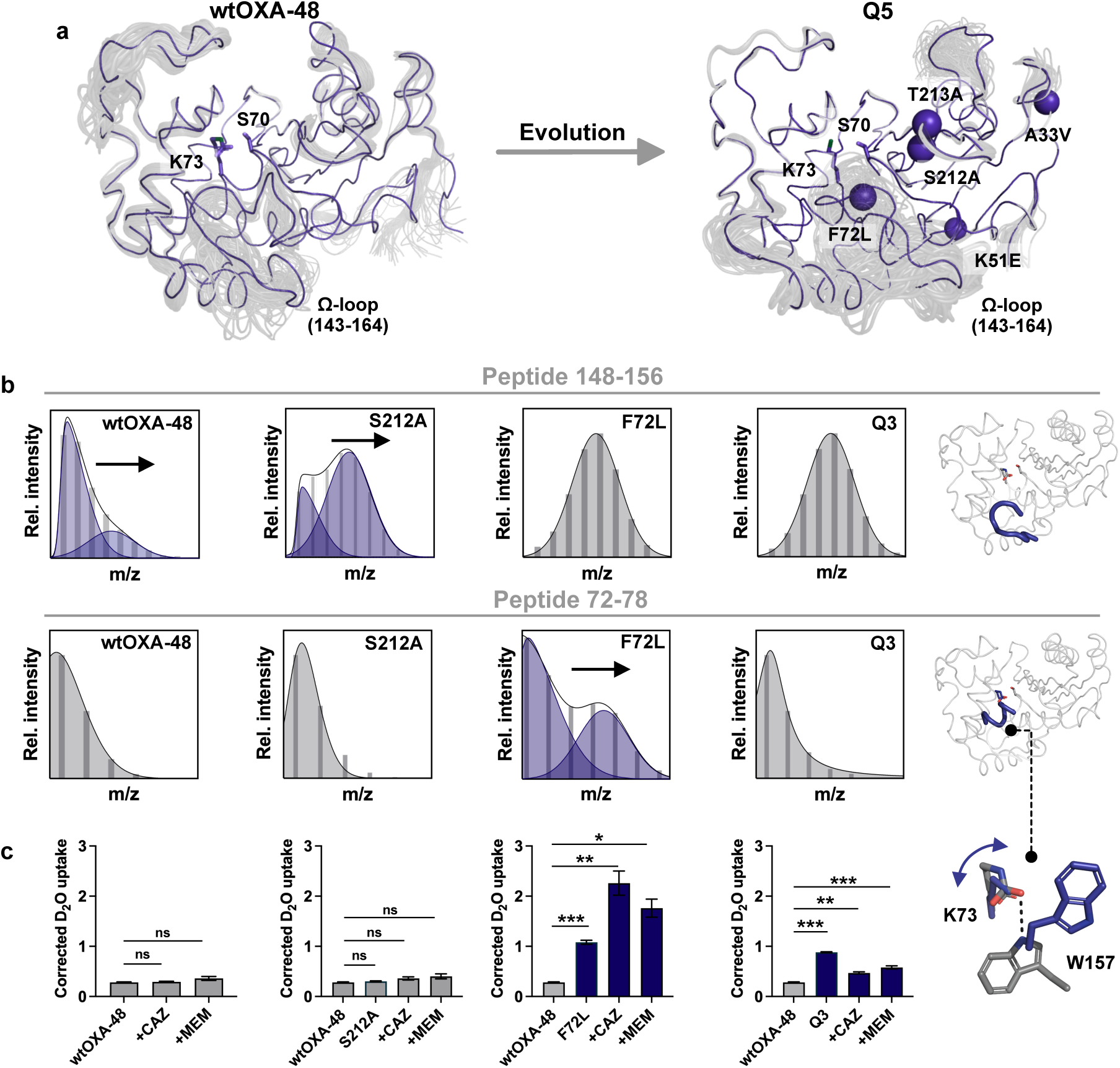
Changes in D2O uptake using HDX-MS compared to apo wtOXA-48. **a.** X-ray crystal structures combined with ensemble refinement of apo wtOXA-48 and Q5 reveal evolutionary changes in the equilibrium conformation and flexibility of the Ω-loop (PDB IDs: 4S2P^26^ and 8PEB^21^; spheres indicate mutations in Q5). **b.** HDX-MS for F72L (10 min HDX) and S212A (10 s HDX) revealed conformational heterogeneity upon substrate exposure, likely reflecting an evolutionary transition between distinct conformational states (arrows). Peptide 72–78 encompasses S70 and K73, whereas peptide 148–156 corresponds to part of the Ω-loop (for all data see Fig. S12). **c.** HDX-MS for peptide 72-78 harbors the general base K73. No significant changes in D2O uptake were detected between wtOXA-48 and S212A in either the apo form or during turnover. In contrast, F72L and Q3 (purple) exhibited significantly increased D₂O uptake under all conditions. Statistical significance was assessed using one-way ANOVA with Welch’s correction, followed by Dunnett’s T3 post hoc test (ns = p > 0.05; * = p < 0.05; ** = p < 0.01; *** = p < 0.001). D2O uptake corrected for back-exchange.^28^ Error bars reflect the standard deviation. For the whole sequence data, see Fig. S13.

To probe the conformational dynamics of the functional sub-states in solution, we analyzed wtOXA-48, S212A, F72L, and Q3 by hydrogen/deuterium exchange mass spectrometry (HDX-MS). Because heterogeneous conformations are expected to produce distinct D_2_O exchange patterns, we hypothesized that HDX-MS could enable characterization of evolutionary changes in conformational sub-states (see Supplementary Text). To that end, we first tested the apo enzymes (Fig. S5). Consistent with increased Ω-loop flexibility, all apo enzymes, except S212A, displayed elevated D_2_O uptake in this region compared to wtOXA-48 (Fig. S5). While the apo enzymes exhibited homogeneous exchange behavior that did not resolve individual sub-states, assays during turnover with ceftazidime and meropenem revealed two distinct populations for S212A and F72L (Fig. 4b, Tab. S3, and Fig. S6 to S12). For F72L, heterogeneity was observed around the catalytic residues S70 and K73, whereas for S212A, two conformational states were observed at the Ω-loop (Fig. 4b and Fig. S12). Notably, wtOXA-48 and Q3 remained homogeneous upon substrate addition, supporting an evolutionary shift in functional sub-states via flexible and heterogeneous evolutionary intermediates (Fig. 2d and Fig. 3c).

Next, we tested whether differences in functional sub-state dynamics could account for the activity trade-off between ceftazidime and meropenem by comparing D_2_O uptake during substrate turnover with apo wtOXA-48 (Fig. S13). F72L and Q3, which exhibited a strong activity trade-off for meropenem, displayed a significant increase in D_2_O uptake in peptide 72–78, close to the nucleophilic Ser70 and including the catalytic base K73 (one-way ANOVA with Welch’s correction, followed by Dunnett’s post hoc test, p < 0.01). This increase was detectable both in the apo forms and under turnover conditions (Fig. 4c). Although substrate exposure also affected D_2_O uptake in other regions, these changes were similar across variants and therefore are unlikely to account for the trade-off (Fig. S13). Accordingly, increased flexibility of the S70/K73 active-site region was specific to F72L and Q3 and absent in wtOXA-48 and S212A. The high catalytic efficiency of wtOXA-48 for meropenem likely relies on a precisely organized active site. Disrupting this organization by increasing flexibility would therefore be expected to reduce turnover. In contrast, the low ceftazidime activity of wtOXA-48 likely reflects misalignment at the active site, in which case increased flexibility may be beneficial by enabling sampling of alternative, catalytically superior conformations (Fig. 4c and Fig. S13).

We further probed the molecular origin of the altered active-site flexibility observed by HDX-MS using molecular dynamics (MD) simulations of wtOXA-48 and Q4 (Fig. S14). Consistent with observations from HDX-MS, MD simulations revealed a significant increase in the flexibility of the catalytic base K73 in Q4 compared to wtOXA-48 across apo, ceftazidime-bound, and meropenem-bound structures (*Δ*RMSF = 0.44, 0.18, and 0.29 Å, respectively; two-tailed t-test: p ≤ 0.01, Tab. S4). In wtOXA-48, the carbamylated K73 forms a hydrogen bond with W157 in the Ω-loop, whereas conformational changes weaken this interaction in the Q4 Ω-loop, resulting in enhanced flexibility of K73 (Fig. 4c, Fig. S15 and S16). This altered interaction between the Ω-loop and the catalytic base provides a plausible mechanistic basis for the evolution of a functional sub-state that favors ceftazidime binding while being less productive for meropenem hydrolysis.

In conclusion, our results suggest that evolution stepwise selected for functional sub-states with distinct conformational flexibility. The change in conformational flexibility around the catalytic serine introduced by F72L likely drives the trade-off with meropenem activity. While this evolutionarily acquired feature facilitates ceftazidime hydrolysis, it appears to impair the chemical step during meropenem hydrolysis.

## DISCUSSION

A central question in enzyme evolution is how mutations reshape conformational landscapes to alter catalytic activity. It remains unclear whether and how evolution can tune the equilibrium between functional sub-states to favor catalytically superior states. In OXA-48, mutations progressively shifted the rate-limiting step from ceftazidime binding to sub-state interconversion rates and ultimately to the chemical step (Fig. 2). Tuning functional sub-state interconversion thereby enriched a more catalytically productive population (Fig. 2). Although Tawfik and colleagues have long argued that evolution of new functions can act by reshaping conformational ensembles,^19^ it has been difficult to probe experimentally how evolution affects functional sub-states, due to challenges in resolving and characterizing these states. While many studies report evolutionary changes in conformational ensembles, such shifts are seldom experimentally assayed to confirm their catalytical relevance during evolution.^6,29–31^ Our work demonstrates how the evolution of functional sub-state dynamics relates directly to catalytic activity. Establishing how general this mechanism is will likely require combining in-depth kinetic analyses with advanced structural techniques such as time-resolved crystallography, HDX-MS, or NMR spectroscopy.

Kinetic analyses with the trade-off substrate meropenem show that optimization for one substrate can shift the functional sub-state balance at the expense of another (Fig. 3). Similarly, recent work on kynureninases demonstrated that divergent conformational ensembles can alter substrate preferences.^32^ Our data underline the crucial role of sub-state evolution in shaping substrate specificity and functional diversification. Accordingly, divergent evolution of conformational flexibility can create substrate incompatibility between different functional sub-states, thereby shaping the specialization and promiscuity of enzymes.

The evolutionary tuning of sub-state interconversion kinetics in OXA-48 carries important implications for enzyme design and engineering. A certain degree of conformational flexibility could be advantageous in early engineering stages, as flexibility may enable access to alternative sub-states with increased catalytic potential. Moreover, flexibility may buffer against inaccuracies introduced during enzyme design by allowing sampling of compensatory conformations.^33^ Notably, many *de novo* designed proteins are highly rigid, which limits the accessible conformational ensemble and may reduce their engineerability compared to natural enzymes. Incorporating conformational flexibility into enzyme design frameworks may therefore enhance both catalytic activity and engineerability.^29,34,35^

Overall, our work demonstrates that kinetic bottlenecks within the conformational landscape can act as direct targets for optimization, enabling evolution to rewire catalytic cycles by accessing distinct functional sub-states. We propose that such evolutionary strategies may be more common than currently appreciated. Uncovering them will likely require detailed kinetics paired with innovative structural approaches. Understanding functional sub-state evolution may ultimately improve the predictability of enzyme evolution and open new routes for rational biocatalyst design.

## METHODS

### General material

Lysogeny broth (LB), LB agar, chloramphenicol, ceftazidime and meropenem were purchased from Sigma-Aldrich (Saint Louis, MO, USA).

### Half-maximal inhibitory concentration (*IC_50_*) susceptibility testing

Susceptibility testing was performed by microbroth dilution assays in 384-well plates (Thermo Fisher Scientific) in two-fold dilution series of either ceftazidime or meropenem, as previously described.^21,36^ In brief, LB broth was inoculated with 10^5^ - 10^6^ CFU/mL and plates were incubated at 37 °C for 20 h. *OD*_600_ was determined using a microtiter plate reader (BioTek Epoch 2, Agilent Technologies). Dose-response curves and the corresponding *IC_50_* values were calculated using GraphPad Prism version 9.3.1.

### Protein expression and purification

Protein expression was performed as described previously.^21^ In brief, *E. coli* BL21-AI cultures harboring OXA-48 variants on a pDEST-17 vector were grown to an *OD*_600_ of 0.4 in LB with 100 mg/L ampicillin at 30 °C, 220 rpm. Expression was induced by adding 0.2% L-arabinose and 1 mM IPTG and cultures were incubated at 15 °C, 225 rpm overnight. Harvested cell pellets were resuspended in buffer AL (50 mM HEPES pH 7.2, 50 mM K_2_SO_4_, 0.1% triton, 1x cOmplete protease inhibitor) and sonicated for 1 h on ice. Samples were centrifuged at 4°C, 15,000 rpm for 10 min and supernatant was purified on HisPur Ni-NTA spin columns (Thermo Fisher Scientific). Proteins were eluted using a 50 mM HEPES pH 7.2, 50 mM K_2_SO_4,_ 250 mM imidazole buffer and concentrated using Amicon Ultra tubes (Merck Millipore, Burlington, MA, USA).

### Steady- and pre-steady state kinetics

Kinetic parameters (*k*_cat_*, K*_M_ and *k*_cat_*/K*_M_) of His-tagged variants were determined at steady-state conditions for ceftazidime (Δξ = −9000 M^−1^ cm^−1^, 260 nm) and meropenem (Δξ = −10940 M^−1^ cm^−1^, 300 nm) by measuring the enzymatic reaction in a stopped flow device (Applied Photophysics) at 25 °C. Kinetics were performed in 0.1 M phosphate buffer pH 7.2 supplemented with 50 mM NaHCO_3_ (Sigma-Aldrich) at a final enzyme concentration of 1 µM. Pre-steady-state kinetics were recorded by monitoring changes in tryptophan fluorescence under identical conditions by exciting at 280 nm and monitoring emission changes at 305 nm using a lower cut-off emission filter and a 1 cm excitation pathlength. Single turnover experiments were performed using 15 µM enzyme and 15 µM meropenem. All data were fitted using GraphPad Prism v 9.3.1. Burst phase kinetics and biphasic curves were fitted to Eq. 4 and Eq. 5, respectively.

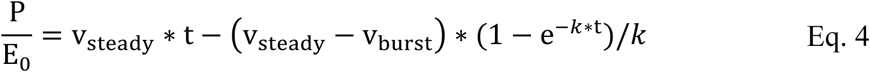

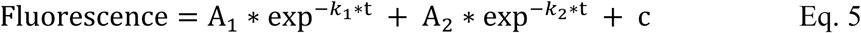

### Crystallization, structural refinement and analysis

Q4 was crystallized as previously described.^21^ Crystals were soaked for 5-10 min in the reservoir supplemented with cryoprotectant using 15% ethylene glycol (Sigma-Aldrich) and piperacillin, and subsequently frozen in liquid nitrogen. Diffraction data were collected on ID30B (Q4-MEM), ESRF, France, at 100 K, wavelength 0.8731 Å,^37^ and the diffraction images were indexed and integrated using XDS.^38^ For data scaling, AIMLESS was used^39^ and an overall high completeness and CC1/2 > 0.5 and a mean intensity above 1.0 in the outer resolution shell was aimed for (Tab. S2). Molecular replacement was performed using chain A of PDB ID 6Q5F^36^ and the program PHENIX v. 1.12^40^. Parts of the model were manually rebuilt using Coot^41^. Average structure refinement and ensemble refinement was performed using PHENIX v. 1.12. PyMOL v. 3.1.3 was used for illustrations (Schrödinger, NY, USA).

### HDX-MS analysis

A detailed method description is part of the Supplementary Information. Briefly, OXA-48 apo variants (1 μM) were dissolved in labelling buffer (50 mM HBS, pD 7.2) in 95.9% D_2_O. H/D exchange was quenched after 10 s, 1 min, 10 min, 1 h, 4 h and 16 h by addition of ice-cold quencher buffer (4 M guanidinium hydrochloride, 0.8 % formic acid) in a 1:1 ratio (final pH of 2.52).^42^ For substrate turnover experiments, enzymes (1 μM) were incubated with ceftazidime or meropenem (850 μM) at different substrate preincubation times (Supplementary Information^43^) and quenched after a 10 s or 10 min H/D exchange. Quenched samples were injected into a Waters HDX Manager in-line with a Xevo G2S ESI Q-TOF (Waters, Milford, MO, USA) mass spectrometer. An Enzymate™ BEH Pepsin Column (Waters) was employed for on-line protein digestion (15 °C and 100 μL/min). Generated peptides were trapped on an Acquity UPLC BEH C18 VanGuard Pre-column (Waters) for 3 min, and eluted on an Acquity UPLC BEH C18 column (Waters) using a linear gradient at 100 μL/min (acetonitrile 2% to 35%, 0.1% formic acid). Maximum deuterated samples for back-exchange correction encompassing mutation sites were prepared.^28^ Peptic fragments generated from online digestion of apo samples were identified with the aid of Protein Lynx Global Server 3.0 (Waters), while D_2_O uptake was analyzed with DynamX 3.0 software (Waters). All DynamX results were manually inspected and subjected to statistical analysis using ANOVA and t-tests (p-value cut-off = 0.05), as implemented in the DECA program.^44^

### MD simulations

The acyl-enzyme structure of wtOXA-48-MEM was built based on the crystal structure (PDB ID: 6P98)^25^. Acyl-enzyme structures of Q4 bound to ceftazidime (Q4-CAZ) and meropenem (Q4-MEM) were built based on the crystal structures of Q5-CAZ (PDB ID: 8PEC)^21^ and Q4-MEM (PDB ID: 9T8V), respectively. To allow for the Ω-loop flexibility observed in structure refinement, the Ω-loop (residues E147-G161) of the Q5 apo form (PDB ID: 8PEB)^21^ was used in Q4 acyl-enzyme structures. The carboxylate group on K73 and deacylating water were added manually to the acyl-enzymes consistent with the reactive acyl-enzyme conformations from previous works^45,46^. Other ionizable residues were modelled in their standard protonation states, with His singly protonated on NE2, consistent with p*K*_a_ prediction and hydrogen bonding. Partial charges and force field parameters of KCX and ceftazidime were taken directly from previous work,^21^ and developed analogously for meropenem. Proteins were treated with the Amber ff14SB force field^47^, with all complexes solvated in a rectangular box of TIP3P water, neutralized with Na^+^ ions. Subsequently, MD simulations were performed as in previous work^21^, with the difference that restraints on KCX and deacylating water were released after the heating stage. For all systems, 32 independent simulations of 40 ns were performed, of which the final 30 ns were used for analysis. See Supplementary Text for further details of setup, parameterisation, simulation and analysis.^48^

## DATA AVAILABILITY

The cryogenic crystal structure was deposited at the Protein Data Bank under the PDB ID: 9T8V (Q4-MEM). Jupyter notebooks and input files required to replicate the MD simulations and analyses of the OXA-48 variants, as well as trajectories and MD snapshots, will be made available on the University of Bristol Research Data Storage Facility (RDSF). All MS raw files will be made available on ProteomeXchange (https://www.proteomexchange.org/) upon publication. Source files will be published alongside the manuscript.

## ACKNOWLEDGEMENTS

DS and CF were supported by the Centre for new antibacterial strategies (CANS) at UiT - The Arctic University of Norway. CF thanks the Norwegian Monitoring Systems for Antibiotic Resistance in Microbes (NORM), the Federation of European Biochemical Societies, the Federation of European Microbiological Societies, and the Northern Norway Regional Health Authority (HNF1722-24). NT thanks the Canadian Institute of Health Research (CIHR) for the project grant (AWD-019305). DW and MWvdK thank the GW4 BIOMED2 DTP for DW’s studentship, grant MR/W006308/1 awarded to the Universities of Bath, Bristol, Cardiff and Exeter from the Medical Research Council (MRC)/UKRI. HAB thanks Swiss National Science Foundation (SNSF) for an Ambizione fellowship (PZ00P3_208691), as well as the Max Planck Society for support. We would like to thank the staff of the ESRF and EMBL Grenoble for assistance and support in using beamline ID30B. MD simulations were conducted using the computational facilities of the Advanced Computing Research Centre, University of Bristol. We are thankful for technical support provided by the Mass Spectrometry Facility of Department of Pharmaceutical and Pharmacological Science at University of Padova.

## AUTHOR CONTRIBUTIONS

DS, HAB and CF performed and analyzed the biochemical assays as well as crystallized and solved structures. AP performed HDX-MS and analyzed the corresponding datasets. KB analyzed and visualized data. DW and MWvdK performed and analyzed MD simulations. CF, NT and HAB designed the study. DS, HAB and CF wrote the manuscript with input from all coauthors.

