## Supplementary Information for "Evolution tunes functional sub-state interconversion to boost enzyme function"

#### Content

1. Supplementary tables 3
   Tab. S1: $IC_{50}$ and *MIC* values of OXA-48 variants selected during directed evolution. 3
   Tab. S2: X-ray data collection and standard refinement statistics. 4
   Tab. S3: HDX incubation time optimization 5
   Tab. S4: K73 side-chain flexibility determined by MD simulations. 6
2. Supplementary figures 7
   Fig. S1: Directed evolution of OXA-48 for ceftazidime hydrolysis. 7
   Fig. S2: Raw traces for binding and turnover kinetics. 8
   Fig. S3: Trade-off development during ceftazidime evolution of OXA-48. 9
   Fig. S4: Structure and ensemble analysis of apo and acyl-enzyme X-ray structures. 10
   Fig. S5: HDX-MS of the apo forms of S212A, F72L, and Q3 compared to wtOXA-48. 11
   Fig. S6: Preincubation time optimization for wtOXA-48 and S212A with ceftazidime. 13
   Fig. S7: Preincubation time optimization for F72L and Q3 with ceftazidime. 14
   Fig. S8: Preincubation time optimization for wtOXA-48 and S212A with meropenem. 15
   Fig. S9: Preincubation time optimization for F72L and Q3 with meropenem. 16

Fig. S10: Effect of ceftazidime and meropenem turnover after 10 s HDX in the respective mutant backgrounds. 17
Fig. S11: Effect of ceftazidime and meropenem turnover after 10 min HDX in the respective mutant backgrounds. 19
Fig. S12: HDX-MS reveals the presence of two populations in F72L and S212A. 21
Fig. S13: Overall $D_2O$ uptake between trade-off and non-trade-off variants. 22
Fig. S14: Comparison of the conformational flexibility between wtOXA-48 and Q4 in apo and acyl-enzyme complexes using MD simulations. 23
Fig. S15: MD simulations display increased side-chain flexibility of the carbamylated K73 24
Fig. S16: Conformational freedom of the K73 side-chain 25
3. Supplementary text 26
3.1 HDX-MS: Extended method 26
3.1.1 Sample preparation 26
3.1.2 HDX-MS 26
3.1.3 Analysis 27
3.2 MD simulations: Extended method 28
4. Supplementary References 30

#### 1. Supplementary tables

**Tab. S1: *$IC_{50}$* and *MIC* values of OXA-48 variants selected during directed evolution.**

| OXA-48 variants | $IC_{50}$ CAZ (mg/L) [a] | *MIC* CAZ (mg/L) [a] | $IC_{50}$ MEM (mg/L) | *MIC* MEM (mg/L) |
|---|---|---|---|---|
| *E. coli* E. cloni | 0.011 ± 0.001 | 0.03 | 0.019 ± 0.001 | 0.06 |
| wtOXA-48 | 0.013 ± 0.002 | 0.03 | 0.300 ± 0.034 | 1 |
| A33V | 0.017 ± 0.004 | 0.03 | 0.297 ± 0.026 | 1 |
| F72L | 0.029 ± 0.006 | 0.12 | 0.025 ± 0.001 | 0.06 |
| S212A | 0.015 ± 0.002 | 0.03 | 0.128 ± 0.009 | 0.5 |
| T213A | 0.012 ± 0.001 | 0.03 | 0.217 ± 0.020 | 1 |
| A33V/F72L | 0.034 ± 0.002 | 0.12 | 0.042 ± 0.002 | 0.125 |
| A33V/S212A | 0.014 ± 0.000 | 0.03 | 0.128 ± 0.006 | 0.25 |
| A33V/T213A | 0.015 ± 0.001 | 0.03 | 0.196 ± 0.014 | 0.5 |
| F72L/S212A | 0.140 ± 0.003 | 0.5 | 0.028 ± 0.001 | 0.06 |
| F72L/T213A | 0.177 ± 0.063 | 1 | 0.036 ± 0.001 | 0.125 |
| S212A/T213A | 0.023 ± 0.005 | 0.06 | 0.140 ± 0.013 | 0.5 |
| A33V/F72L/S212A | 0.148 ± 0.042 | 0.5 | 0.031 ± 0.001 | 0.06 |
| A33V/F72L/T213A | 0.141 ± 0.028 | 0.5 | 0.027 ± 0.001 | 0.06 |
| A33V/S212A/T213A | 0.020 ± 0.003 | 0.06 | 0.061 ± 0.003 | 0.125 |
| F72L/S212A/T213A (Q3) | 0.389 ± 0.025 | 1 | 0.025 ± 0.001 | 0.06 |
| A33V/F72L/S212A/T213A (Q4) | 0.513 ± 0.093 | 1 | 0.034 ± 0.002 | 0.06 |
| A33V/K51E/F72L/S212A/T213A (Q5) | 0.556 ± 0.052 | 1 | 0.035 ± 0.001 | 0.06 |

Errors are reported as the standard error of the mean. The average is calculated based on at least 3 biological replicates.

[a] CAZ $IC_{50}$ and *MIC* values were obtained from[1]

57 **Tab. S2: X-ray data collection and standard refinement statistics.**

| | Q4-MEM |
|---|---|
| PBP | 9T8V |
| Beamline | ID30B, ESRF |
| Wavelength (Å) | 0.8731 |
| Resolution range (Å) | 45.2-2.1 (2.175-2.1) |
| Space group | P $2_1$ $2_1$ $2_1$ |
| Unit cell: a,b,c (Å) | 64.7, 82.2, 98.8 |
| α, β, γ (°) | 90, 90, 90 |
| Total reflections | 62828 (6204) |
| Unique reflections | 31417 (3102) |
| Multiplicity | 2 |
| Completeness (%) | 99.7 (99.4) |
| Mean I/sigma(I) | 17.8 (1.8) |
| Overall *B*-factor from Wilson plot (Å$^2$) | 50 |
| $R_{merge}$ | 0.01776 (0.4117) |
| $R_{measured}$ | 0.02512 (0.5822) |
| $R_{pim}$ | 0.01776 (0.4117) |
| $CC_{1/2}$ | 1 (0.871) |
| Resolution range (Å) | 25 - 2.1 |
| Reflections used in refinement | 31345 (3090) |
| Reflections used for R-free | 1582 (150) |
| Final $R_{work}$ | 0.2080 (0.3258) |
| Final $R_{free}$ | 0.2811 (0.3805) |
| No. of non-hydrogen atoms | |
| -macromolecules | 3850 |
| -ligands | 62 |
| -solvent | 64 |
| R.m.s. deviations | |
| -bonds (Å) | 0.008 |
| -angles (°) | 0.90 |
| Ramachandran plot | |
| -Favoured (%) | 95.2 |
| -Allowed (%) | 3.5 |
| -Outliers (%) | 1.3 |
| Average B-factor (Å$^2$) | 60 |
| -macromolecules (Å$^2$) | 60 |
| -ligands (Å$^2$) | 73 |
| -solvent (Å$^2$) | 54 |

58 Statistics for the highest-resolution shell are shown in parentheses.

#### Tab. S3: HDX incubation time optimization

| States | Preincubation times (min)[c] 0[a] | 1 | 10 | 30 | 60 | HDX[d] time | Observed changes in $D_2O$ uptake[b] |
|---|---|---|---|---|---|---|---|
| wtOXA-48 + CAZ | | | x | x | x | 10 s | Only minor differences were observable after 60 min preincubation and mainly in the Ω-loop (Fig. S6). |
| wtOXA-48 + CAZ | | | | x | x | 10 min | |
| wtOXA-48 + MEM | x | | | | | 10 s | Substantial changes in $D_2O$ uptake were detectable without preincubation, including the Ω-loop and the α4-α5 and α6 helices (Fig. S8). |
| wtOXA-48 + MEM | x | | | | | 10 min | |
| S212A + CAZ | | | x | x | x | 10 s | Substantial $D_2O$ uptake changes detectable after both a 10 and 30 min preincubation. Minimal differences in uptake were observed between 30 and 60 min (Fig. S6). Major changes included the Ω-loop and the α4-α5 and α6 helices. |
| S212A + CAZ | | | | x | | 10 min | |
| S212A + MEM | x | x | | | | 10 s | Substantial differences in $D_2O$ uptake were observed after both preincubation times (Fig. S8). Major changes were observed in α6-helix and parts of the α4-α5-helix. |
| S212A + MEM | x | x | | | | 10 min | |
| F72L + CAZ | | x | x | | | 10 s | Substantial differences in $D_2O$ uptake were observed after 10 min preincubation. Smaller effects are detected after a 1 min preincubation, mainly around the catalytic residues S70 and K73 (Fig. S7). |
| F72L + CAZ | x | | x | | | 10 min | |
| F72L + MEM | | x | x | | | 10 s | After 1 min of preincubation, substantial changes in $D_2O$ uptake were detected (Fig. S9). Effect magnitude decreased at 10 min preincubation time. The main effect was an increase in $D_2O$ uptake around the catalytic residues S70 and K73. |
| F72L + MEM | x | x | x | | | 10 min | |
| Q3 + CAZ | x | x | x | | | 10 s | Substantial differences in $D_2O$ uptake are visible after both a 1 and 10 min preincubation (Fig. S7). Main differences were around the α6-helix and the α4-α5-helix. |
| Q3 + CAZ | | x | | | | 10 min | |
| Q3 + MEM | | x | x | | | 10 s | Substantial $D_2O$ uptake is detectable with minor differences across 0, 1, or 10 min preincubation. (Fig. S9) |
| Q3 + MEM | x | x | x | | | 10 min | |

[a] After 0 min indicates that samples were mixed and injected immediately.

[b] Substantial changes in $D_2O$ uptake were defined as differences > |± 0.3| Da and |± 4%| relative to the corresponding apo variants. Except for wtOXA-48, all variants exhibited less $D_2O$ exchange at the binding site, consistent with substrate occupancy of the active site. These changes are not mentioned in Tab. S4. It is important to note that under substrate turnover, the signal intensity of residues 65 to 71 was reduced for all variants except for wtOXA-48. This is a direct consequence of the covalent modification of S70 that occurs during substrate turnover.[2]

[c] Preincubation times with either ceftazidime (CAZ) or meropenem (MEM). Preincubation times used for downstream analysis are indicated in blue

[d] Hydrogen/Deuterium exchange (HDX) times.

**Tab. S4: K73 side-chain flexibility determined by MD simulations.**

| System[a] | wtOXA48 RMSF (Å) | Q4 RMSF (Å) | p value[b] |
|---|---|---|---|
| Apo states | 0.59 ± 0.09 | 1.03 ± 0.25 | $1.48 \times 10^{-11}$ |
| Ceftazidime-bound | 0.50 ± 0.02 | 0.68 ± 0.29 | $1.29 \times 10^{-3}$ |
| Meropenem-bound | 0.50 ± 0.01 | 0.80 ± 0.33 | $1.59 \times 10^{-5}$ |

[a] Root-mean square fluctuation (RMSF) values of the carbamylated lysine side-chain heavy atoms were calculated by aligning each trajectory to the average coordinate set based on the carbamylated lysine backbone heavy atoms.

[b] An independent two-sample, two-tailed t-test was performed to compare the RMSF values between wtOXA-48 and Q4 (n = 32).

Errors represent the standard deviation.

#### 2. Supplementary figures

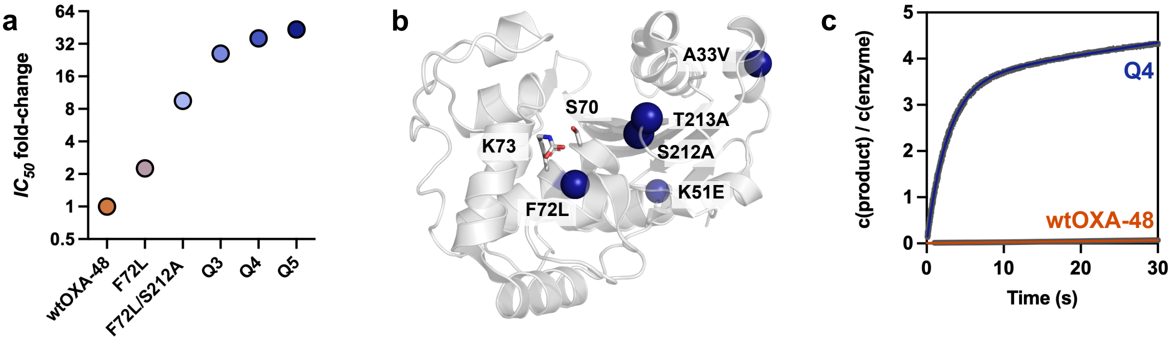


**Fig. S1: Directed evolution of OXA-48 for ceftazidime hydrolysis.**
**a.** Determined $IC_{50}$ values along the evolutionary trajectory from wtOXA-48 to Q5 (F72L→S212A→T213A→A33V→K51E) show a gradual increase in ceftazidime resistance, reaching up to a 43-fold improvement. **b.** Mutations (spheres) acquired during evolution are mapped onto the OXA-48 structure (PDB ID: 4S2P), along with key catalytic residues S70 and K73. **c.** Kinetic analysis shows increased ceftazidime turnover for Q4 compared to wtOXA-48, introducing a super-stoichiometric burst phase. Figure adapted from[1].

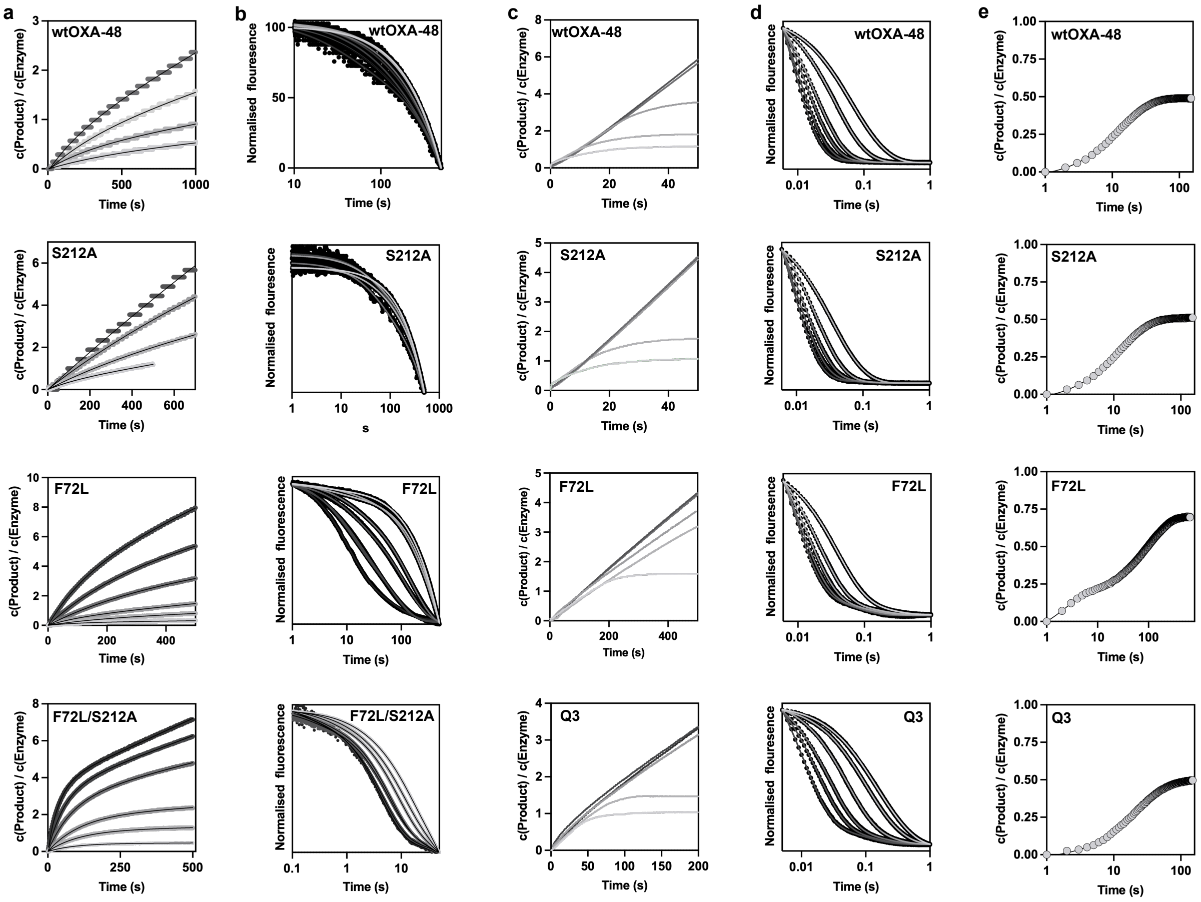


**Fig. S2: Raw traces for binding and turnover kinetics.**

**a.** Ceftazidime turnover progress curves for substrate concentrations ranging from 5 to 400 µM. **b.** Ceftazidime binding kinetics monitored by tryptophan fluorescence for concentrations ranging from 50 to 1200 µM. **c.** Meropenem turnover progress curves for substrate concentrations ranging from 1.25 to 25 µM. Plateauing of the curves does not reflect super-stoichiometric burst behaviour but substrate depletion. **d.** Meropenem binding kinetics monitored by tryptophan fluorescence for concentrations ranging from 1.25 to 25 µM. **e.** Single turnover experiments with meropenem (c(enzyme) = 15 µM, c(meropenem) = 15 µM). Data analysis is shown in Fig. 2 and 3 as well as Tab. 1 and 2. All experiments were performed at least in duplicates.

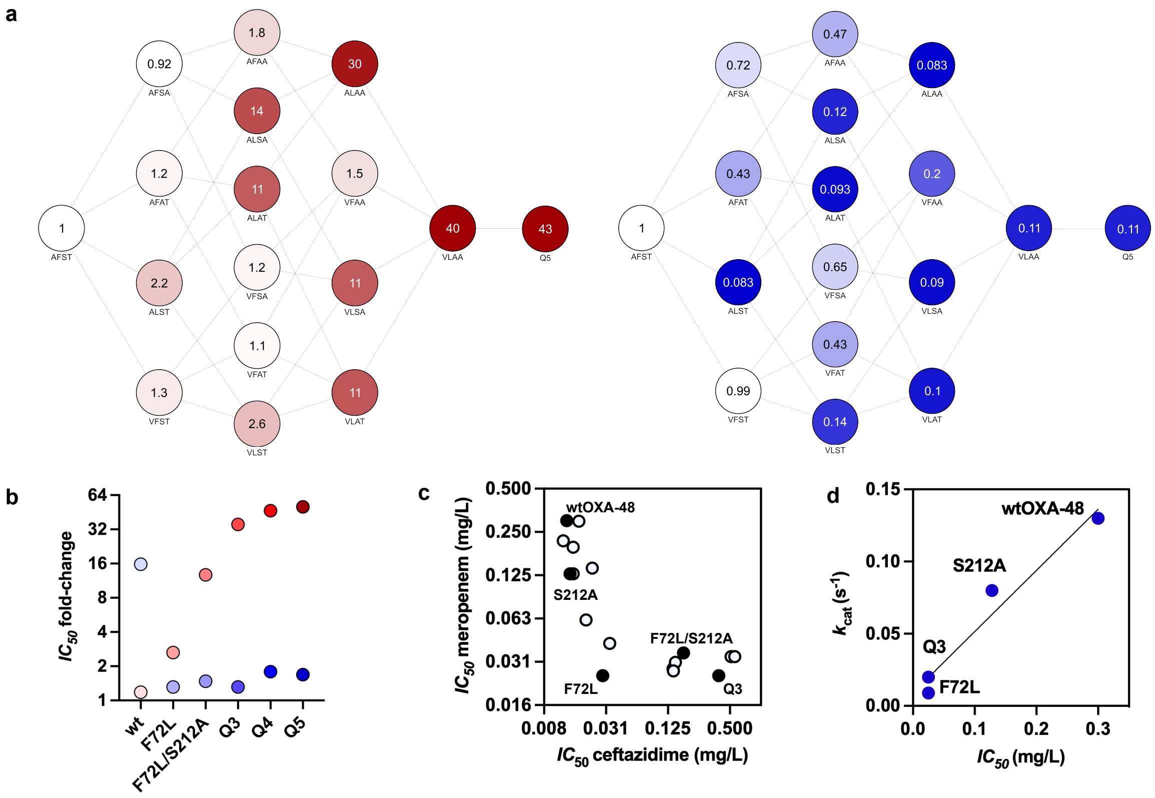


**Fig. S3: Trade-off development during ceftazidime evolution of OXA-48.**

**a.** Adaptive fitness landscapes of the four accumulated mutations (A33V, F72L, S212A, and T213A) and the endpoint mutant Q5 (A33V/K51E/ F72L/S212A/T213A) for ceftazidime (red) and meropenem (blue). $IC_{50}$ values for all variants are listed in Tab. S1. **b.** $IC_{50}$ values along the evolutionary trajectory for ceftazidime resistance (red) showed that the addition of F72L substantially reduced meropenem resistance (blue). **c.** The $IC_{50}$ values of the adaptive landscapes show a strong trade-off between OXA-48-mediated resistance to ceftazidime and meropenem. Filled dots indicate variants analysed in detail in this study. **d.** The strong correlation between meropenem $IC_{50}$ and $k_{cat}$ values indicates that the observed trade-off is driven by changes in the enzyme turnover number (Pearson correlation, $R^2 = 0.95$, p = 0.02).

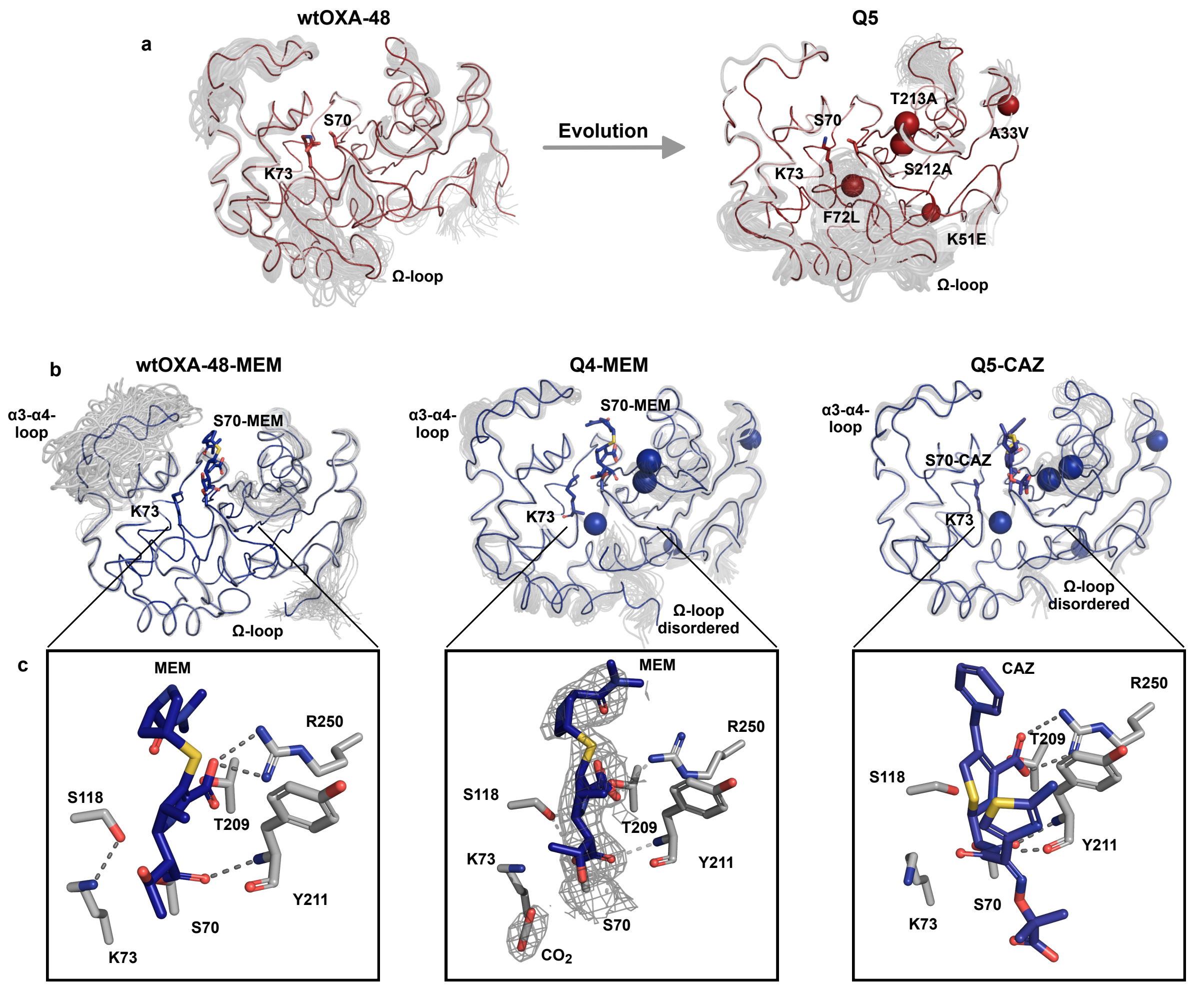


**Fig. S4: Structure and ensemble analysis of apo and acyl-enzyme X-ray structures.**

Structural analysis of the evolutionary trajectory of wtOXA-48 (PDB ID: 4S2P)[3] to Q5 (PDB ID: 8PEB)[1] for increased ceftazidime resistance. Chain A is shown for all structures. Average structures are shown in colour, and ensembles from ensemble refinement are shown in grey. The locations of mutations introduced during evolution are highlighted as spheres. **a.** Apo structures reveal that the conformation of the Ω-loop changed during evolution. Moreover, the flexibility of the Ω-loop increased, which was also observed in MD simulations (Fig. S14).[1] **b.** Global effect of substrate binding on the average structure and ensembles. The acyl enzyme structures of wtOXA-48 with meropenem (wtOXA-48-MEM, PDB: 6P98)[4], Q4 with meropenem (Q4-MEM; PDB ID: 9T8V), and Q5 with ceftazidime (Q5-CAZ; PDB ID:8PEC)[1] are analysed. Electron density map for meropenem (MEM) in Q4 is shown as 2Fo-Fc maps at 1σ. In wtOXA-48, no substantial increases in ensemble conformations, except for the α3-α4-loop, were observed. The Ω-loop is in the typical closed conformation.[5] In the ceftazidime-evolved variants (Q4/Q5), the Ω-loops could not be refined in either of the acyl-enzyme complexes due to a lack of electron density, indicating highly dynamic loop motions upon substrate binding. **c.** Close-up view of the substrate-bound active site. Dashed lines indicate polar interactions. R250 likely acts as an anchor, stabilizing the negative charge on the carboxylic group in β-lactams. In both wtOXA-48-MEM and Q5-CAZ, R250 is optimally positioned. In Q4-MEM, R250 is tilted away from the substrate, leading to less favourable interactions.

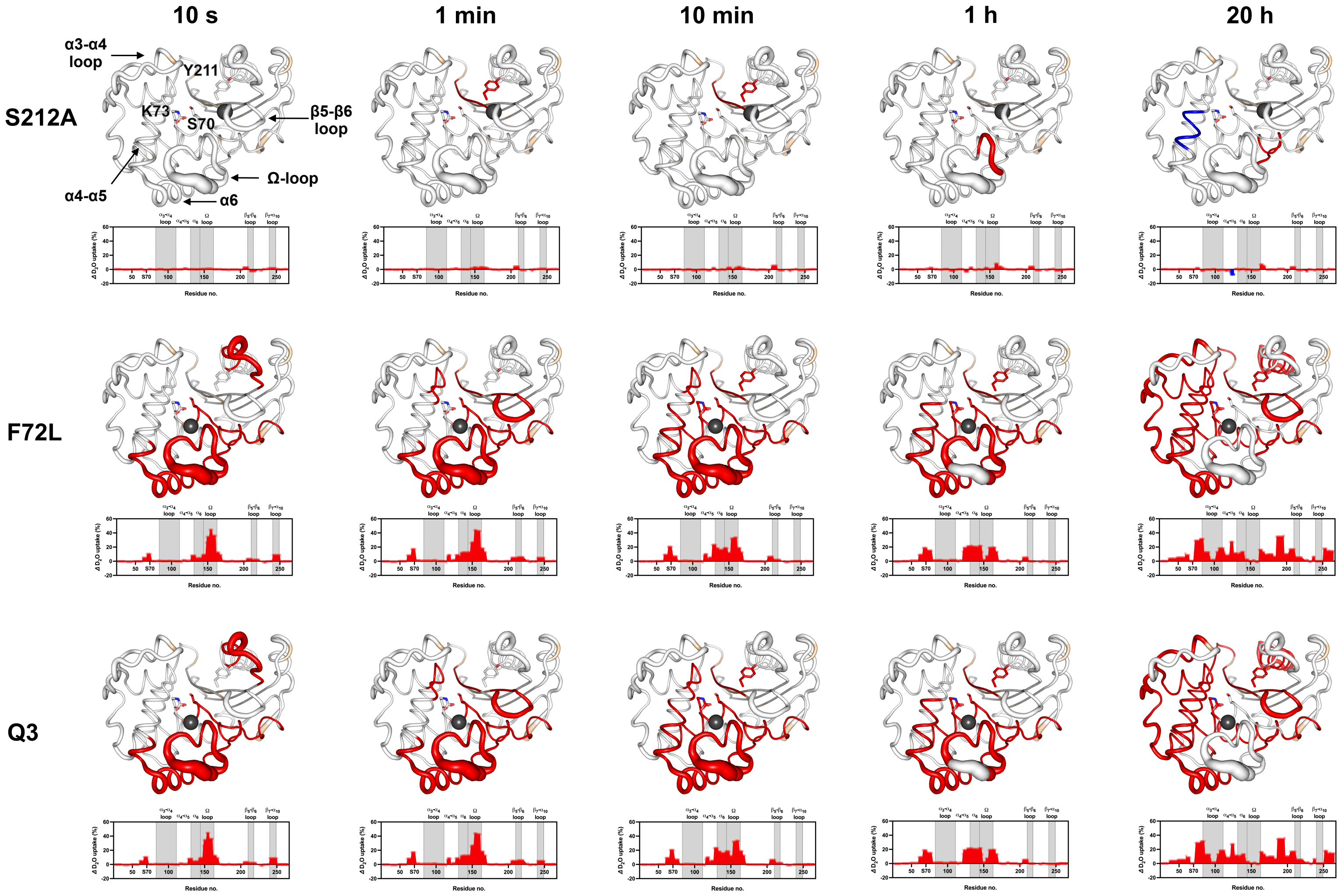


**Fig. S5: HDX-MS of the apo forms of S212A, F72L, and Q3 compared to wtOXA-48.**

Changes in $D_2O$ uptake for S212A, F72L, and Q3 were determined after 10 s, 1 min, 10 min, 1 h, and 16 h of incubation in $D_2O$. $D_2O$ uptake is shown compared to wtOXA-48 at the corresponding time points. Black spheres display the location of the mutations (F72L, S212A, and T213A). Substantial (>4%) increases (red) and decreases (blue) in

$D_2O$ uptake are plotted on the structures. Apo S212A displayed $D_2O$ uptake similar to wtOXA-48 (even after 16 h incubation), indicating that the mutation did not substantially change the conformational flexibility. In contrast, compared to wtOXA-48, both apo F72L and Q3 displayed significantly increased $D_2O$ uptake within the Ω-loop (one-way ANOVA with Welch's correction, followed by Dunnett's T3 post hoc, $p < 0.001$ at 10 s HDX for peptide 148-156 in the Ω-loop), which reached full deuteration at 10s already indicating that in both F72L and Q3 this region becomes fully exposed to $D_2O$. A significant increase in conformational flexibility around the active-site serine (S70) and catalytic base (K73) for both mutants was detectable after 10 min of HDX (one-way ANOVA with Welch's correction followed by Dunnett's T3 post hoc test; $p = 0.007$ and $p = 0.003$ at 10 min for peptide 72–78 in F72L and Q3, respectively, Fig. 4c). Other regions that displayed increased $D_2O$ uptake were: the α6-helix and the β7-α10 loop.

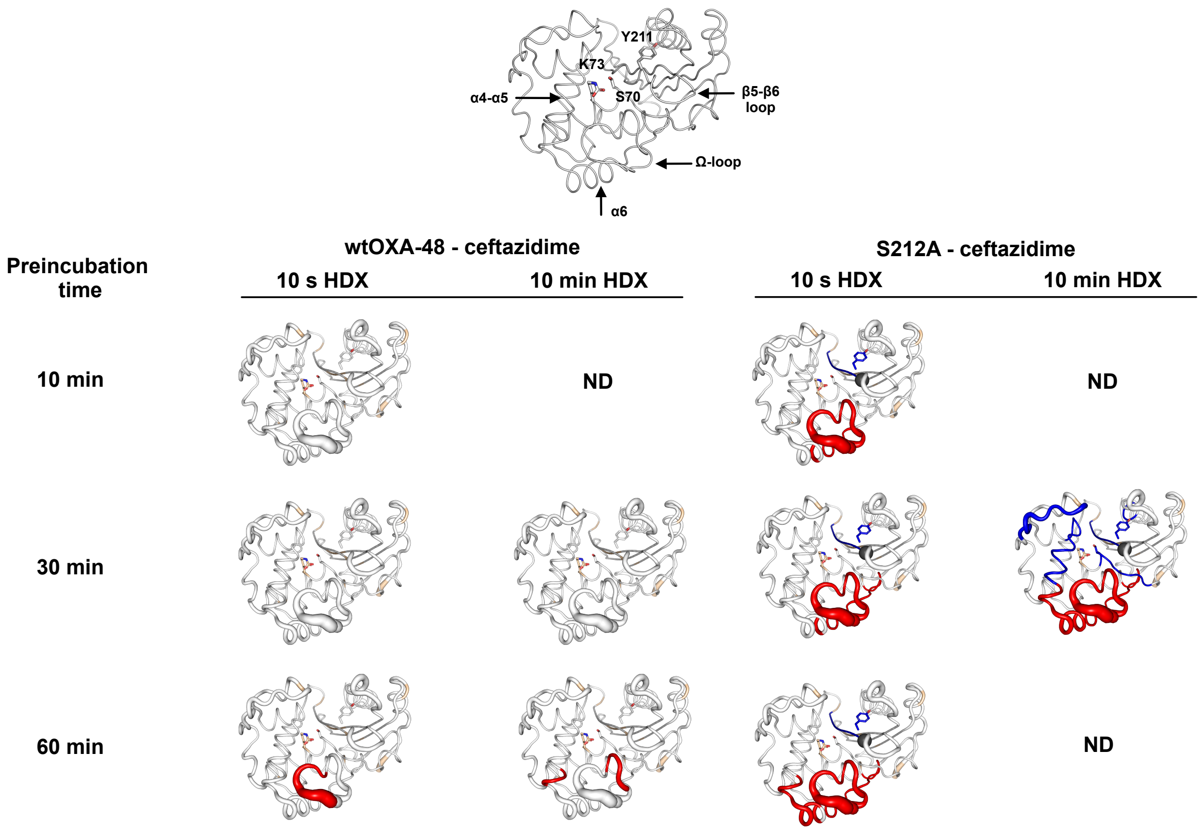


**Fig. S6: Preincubation time optimization for wtOXA-48 and S212A with ceftazidime.** Due to the low $k_{cat}$ of wtOXA-48 and S212A, preincubation times with ceftazidime were probed between 10, 30 and 60 min (Tab. 1). $D_2O$ exchange was probed after 10s and 10 min (HDX). Changes in $D_2O$ uptake greater than |± 4| % are shown on the structures. HDX was recorded with at least two replicates. Changes in $D_2O$ uptake are shown compared to the corresponding apo variant for each preincubation time. Black spheres display the location of the corresponding mutations (F72L, S212A, and T213A). Even after 60 min of preincubation, only marginal changes in $D_2O$ uptake were observed for wtOXA-48. These changes were mainly within the Ω-loop.

The introduction of S212A into wtOXA-48 generally did not affect the $D_2O$ uptake in the apo forms (Fig. S5). In contrast, changes in $D_2O$ in S212A were detectable in the Ω-loop in addition to the α4-α5 and α6 helices during ceftazidime turnover, compared to apo S212A. $D_2O$ uptake was substantially (>4%) higher already after 10 min of preincubation time and 10 s HDX, compared to the apo form of S212A. In addition, a >4 % protection from $D_2O$ of the oxyanion hole (including Y211) was observed, indicating the presence of the substrate in the binding pocket. Prolonged preincubation times of up to 60 min did not lead to additional substantial changes in $D_2O$ at 10 s HDX. Substrate-induced protection became even more pronounced after 10 min of HDX and 30 min preincubation with ceftazidime. 60 min (wtOXA-48) and 30 min (S212A) preincubation times were included into downstream analysis (Fig. S10 and Fig. S11). ND = Not determined.

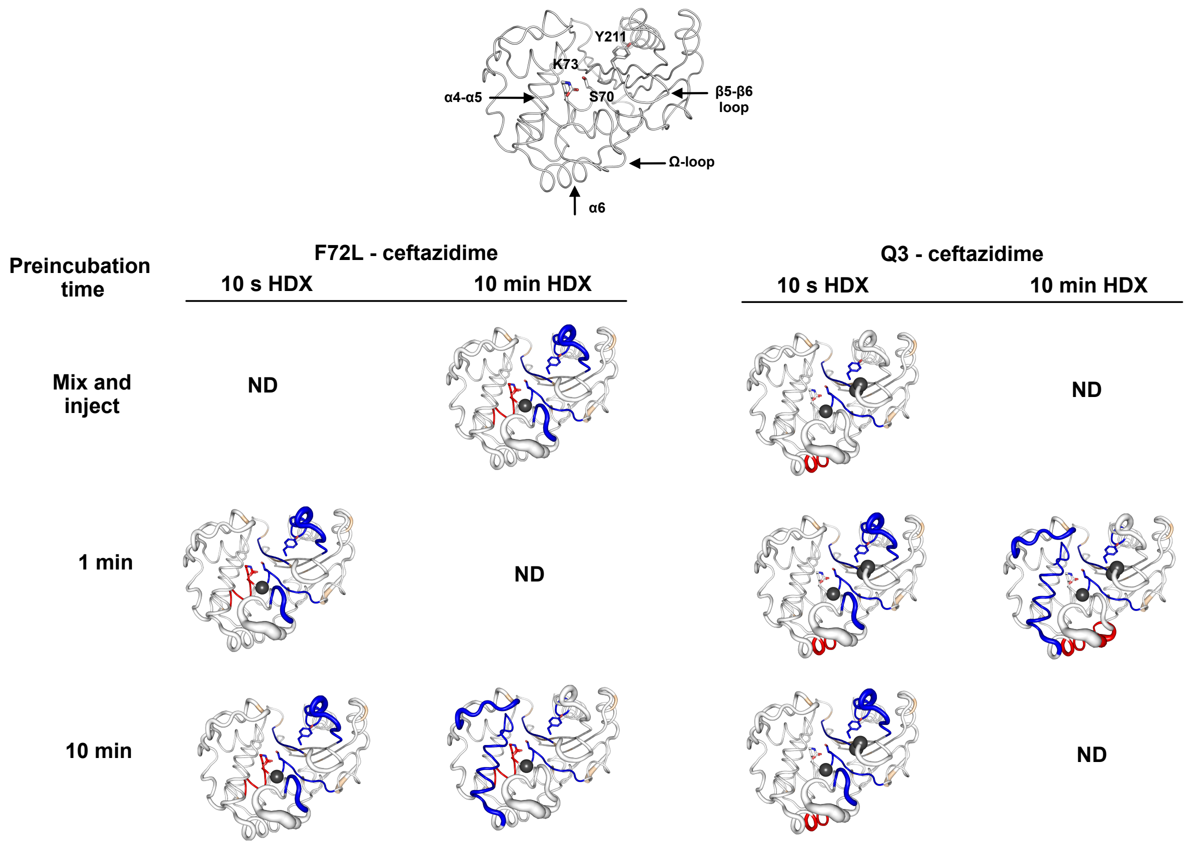


**Fig. S7: Preincubation time optimization for F72L and Q3 with ceftazidime.**

Due to the increased $k_{cat}$ of F72L and Q3 compared to their ancestral variants (Tab. 1), shorter substrate preincubation times of 10 s, 1 min, and 10 min with ceftazidime were explored. $D_2O$ exchange was probed after 10s and 10 min (HDX). Changes in $D_2O$ uptake greater than |± 4| % are shown on the structures. HDX was recorded with at least two replicates. Changes in $D_2O$ uptake are shown compared to the corresponding apo variant for each preincubation time. Black spheres display the location of the corresponding mutations (F72L, S212A, and T213A). While mutations in F72L and Q3 generally increased the $D_2O$ uptake of regions around the active site, including the region around the active site serine, the α4-α5-helix, the α6-helix, and the Ω-loop (Fig. S5), $D_2O$ uptake was reduced for most of these regions under turnover conditions compared to the apo state of the mutants.

Some of this protection against $D_2O$ exchange is around the ceftazidime binding pocket (around Y211) and is therefore indicative of the presence of the substrate, other regions such as the α6-helix and parts of the α4-α5-helix (F72L: 10 min preincubation and 10 min HDX; Q3: 1 min preincubation and 10 min HDX) are distant from the binding pocket which may be indicative of conformational changes occurring during turnover. In addition, in all F72L conditions, the region around the active site S70, including the catalytic base K73, displayed even higher $D_2O$ exposure (> 4%) compared to apo F72L (Fig. S5). 10 min (F72L) and 1 min (Q3) preincubation times were included in the downstream analysis (Fig. S10 and Fig. S11). ND = Not determined.

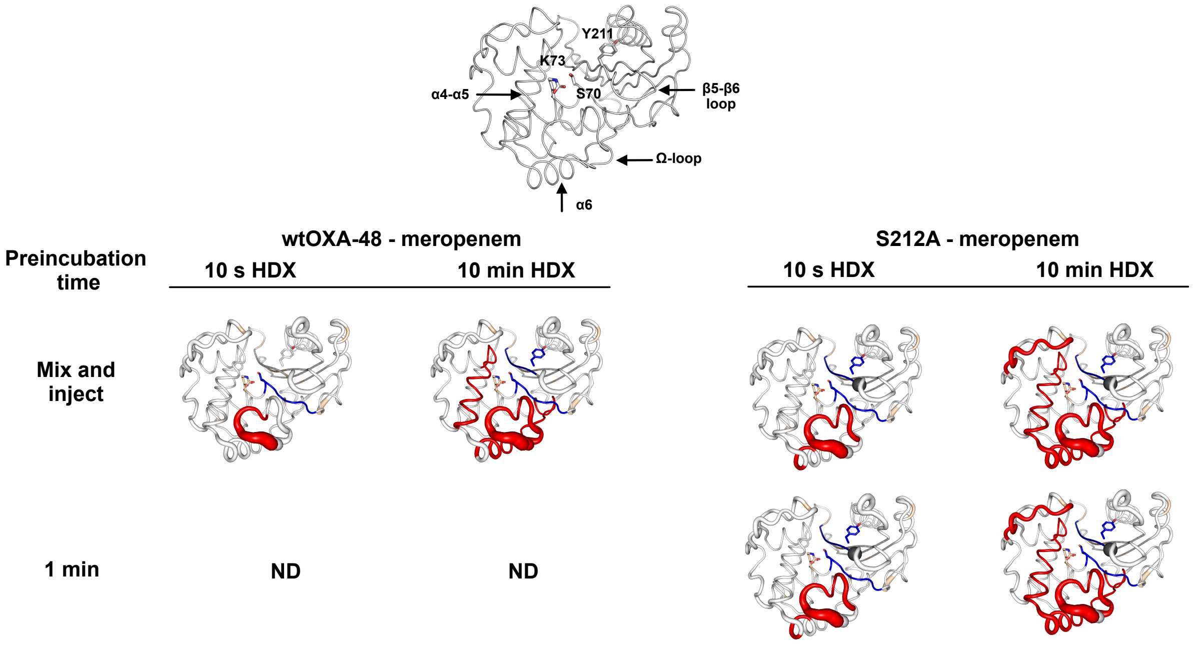


**Fig. S8: Preincubation time optimization for wtOXA-48 and S212A with meropenem.** Due to their efficient meropenem hydrolysis, short preincubation times (mix and inject and 1 min) were tested for wtOXA-48 and S212A. Changes in $D_2O$ uptake greater than |± 4| % are shown on the structures. In addition, 10s and 10 min HDX were explored based on at least two replicates. Changes in $D_2O$ uptake are shown compared to the corresponding apo variant for each preincubation time. Black spheres display the location of the corresponding mutations (F72L, S212A, and T213A).

All conditions displayed protection around meropenem binding pocket (around Y211), indicative of the presence of the substrate in the active site. In addition, for both wtOXA-48 and S212A, meropenem hydrolysis increased the $D_2O$ exposure of the Ω-loop (10 s HDX) in addition to the α4-α5 and α6 helices (10 min HDX) compared to the apo forms of the mutants. Prolonged (1 min) preincubation time in S212A did not alter the changes in $D_2O$ uptake, results obtained from mix and inject were therefore used for both wtOXA-48 and S212A for further analysis (Fig. S10 and Fig. S11).

ND = Not determined, HDX= Hydrogen/deuterium exchange

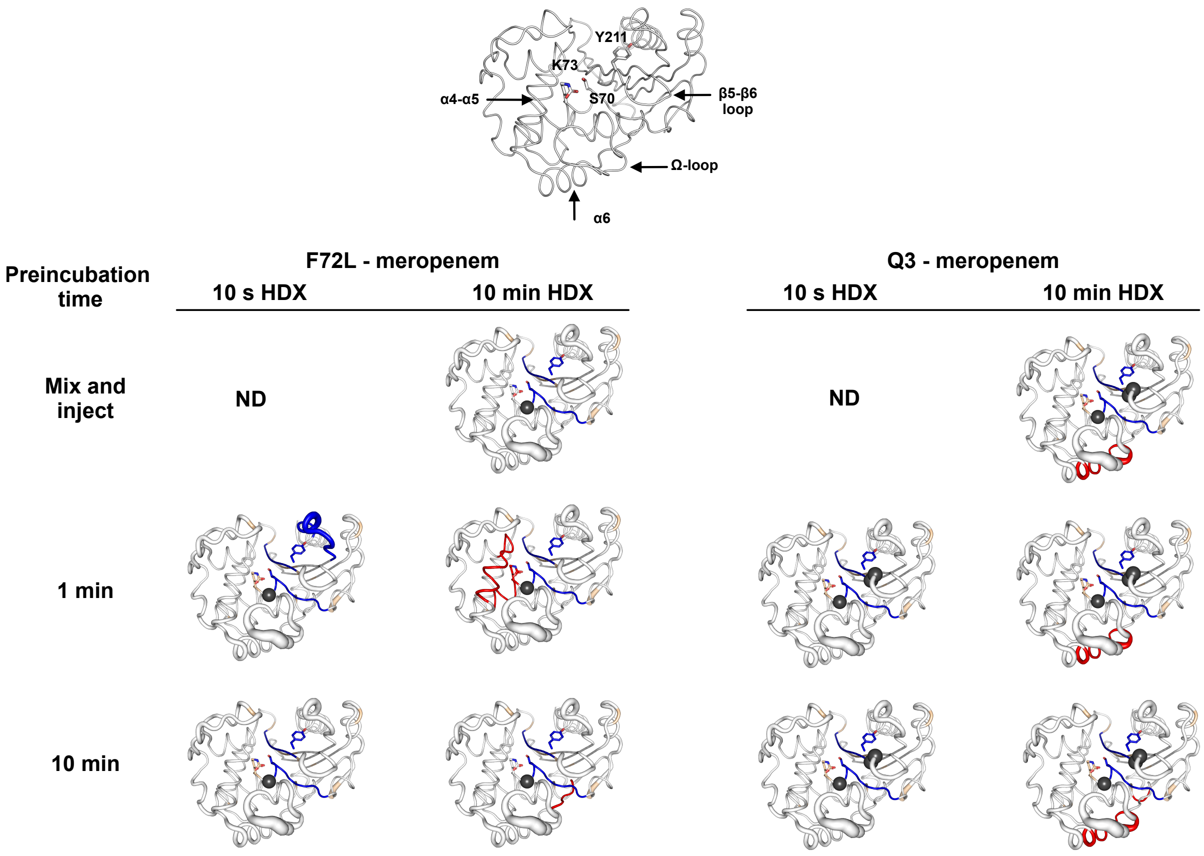


**Fig. S9: Preincubation time optimization for F72L and Q3 with meropenem.**

Due to their lower $k_{cat}$, compared to wtOXA-48, longer preincubation times with meropenem were tested for F72L and Q3: mix and inject, 1 min and 10 min. Changes in $D_2O$ uptake greater than |± 4| % are shown on the structures. In addition, 10s and 10 min HDX were explored based on at least two replicates. Changes in $D_2O$ uptake are shown compared to the corresponding apo variant for each preincubation time. Black spheres display the location of the corresponding mutations (F72L, S212A, and T213A).

Changes in $D_2O$ uptake were compared to the corresponding apo forms of the corresponding variants for each preincubation time. All conditions displayed protection around meropenem binding pocket (around Y211), indicative of the presence of the substrate in the active site. After 10 min with substrate preincubation, a reduction in conformational effects was observed compared to 1 min (F72L). After 10 min of HDX and 1 min preincubation with meropenem, F72L displayed even higher $D_2O$ uptake (> 4%) around the catalytic residues S70 and K73 compared to the apo form of F72L (Fig. S5). 1 min preincubation with meropenem was used for both F72L and Q3 for further analysis (Fig. S10 and Fig. S11).

ND = Not determined, HDX= Hydrogen/deuterium exchange

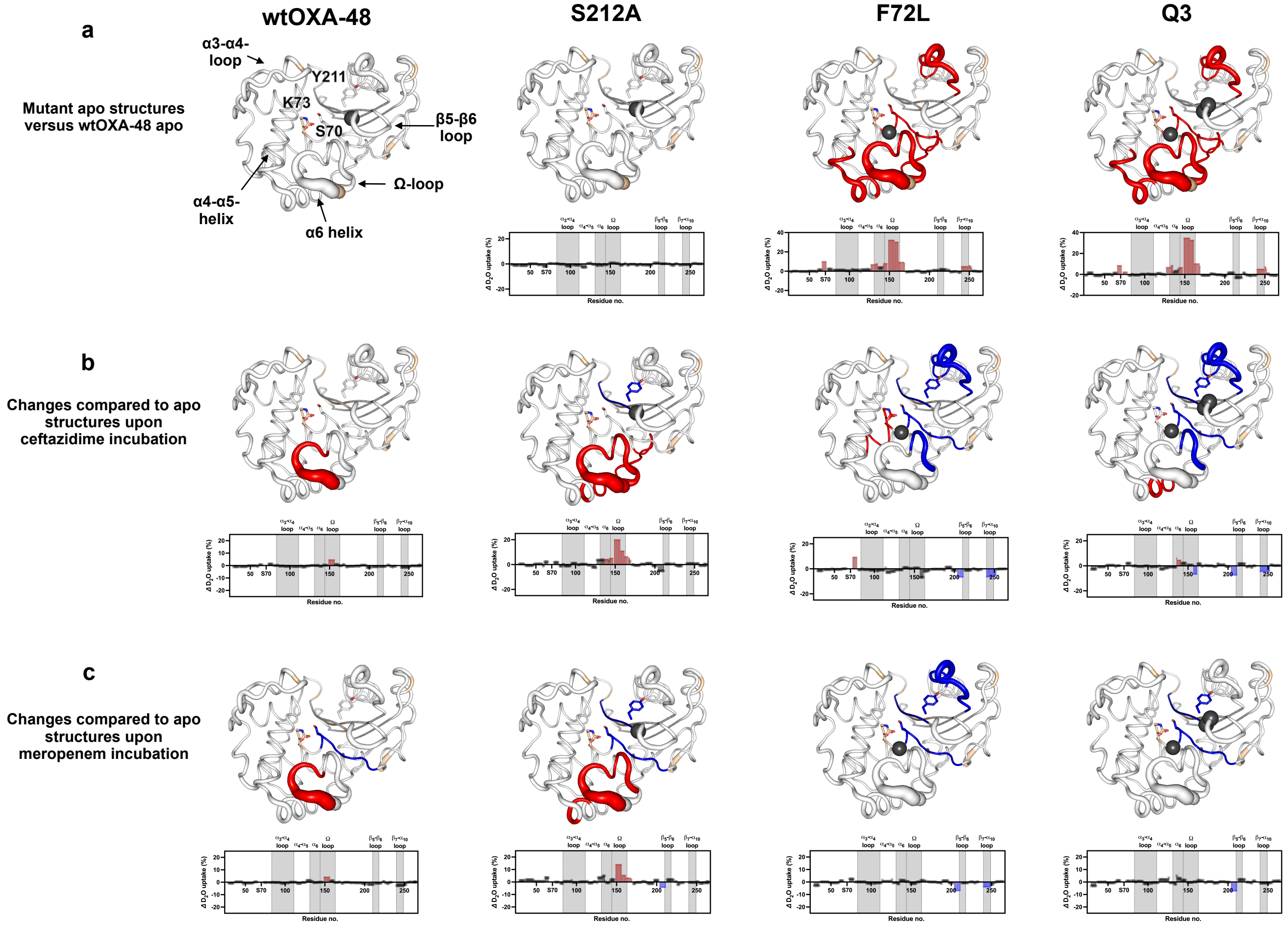


Fig. S10: Effect of ceftazidime and meropenem turnover after 10 s HDX in the respective mutant backgrounds.

Changes in $D_2O$ uptake after 10 s HDX for wtOXA-48, S212A, F72L, and Q3 using HDX-MS. For comparison, changes in $D_2O$ uptake induced by mutations are shown in panel a. Panel b (ceftazidime) and c (meropenem) describe changes during substrate turnover compared to changes inflicted by the mutants (panel a). Preincubation times were optimised based on kinetic parameters (Tab. 1 and Tab. 2) and results summarized in Tab. S3. For ceftazidime, the following substrate preincubation times were used: wtOXA-48 (60 min) , S212A (30 min), F72L (10 min), and Q3 (1 min). For meropenem, the following substrate preincubation times were used: wtOXA-48 (mix and inject) , S212A (mix and inject), F72L (1 min) and Q3 (1 min). For preincubation time optimization see Tab. S3. The relative increase (red) and decrease (blue) in $D_2O$ uptake (>|± 4|%) after 10 min HDX are mapped onto the structures. Black spheres display the location of the corresponding mutations (F72L, S212A, and T213A).

**a.** Panel a shows the effect of mutations on the $D_2O$ uptake compared to apo wtOXA-48. These variants were treated identically to the turnover experiments meaning that they were incubated for the same time frame but without substrate. No substantial changes were observed in the apo form of S212A compared to wtOXA-48. Apo forms of F72L and Q3 experienced a substantial increase in $D_2O$ uptake (15-30%) in the α4-α5 and α6 helices, the **Ω**-loop (peptide 148-156) and around the active site serine (peptides 72-78). Generally these results are in good agreement with our apo HDX experiments without preincubation time (Fig. S5)

**b.** Ceftazidime exposure resulted in limited changes in wtOXA-48, but induced up to 15% increased uptake within the **Ω**-loop and α6-helix of S212A. Decreased $D_2O$ uptake relative to their apo forms in S212A, F72L and Q3 was observed in the α4-α5 and α6 helices and oxyanion hole peptide (with Y211), indicative of substrate binding in the active site. F72L and Q3 under ceftazidime turnover maintained enhanced $D_2O$ uptake compared to their apo forms in the α6-helix and **Ω**-loop, even showing further 10% increased uptake in the α6-helix of Q3. In addition, the region upstream and downstream of the active site serine S70 experienced an up to 15% increase in $D_2O$ exposure in F72L under ceftazidime turnover. While this region rigidified by ca. 4% in the ceftazidime exposed Q3 variants, it remained substantially higher than in the wtOXA-48 and S212A (Fig. 4c).

**c.** Meropenem exposure to wtOXA-48 and S212A led to enhanced $D_2O$ uptake in the α6-helix and **Ω**-loop, resembling the apo forms of F72L and Q3. Although changes in F72L and Q3 were less pronounced, a distinguishing feature of these variants was, similar to ceftazidime, the increased $D_2O$ uptake related to the region upstream and downstream of the active site serine S70 (peptides 72-78). This suggests that enhanced uptake within the active site region may relate to improved ceftazidime hydrolysis, while potentially compromising meropenem hydrolysis.

HDX= Hydrogen/deuterium exchange

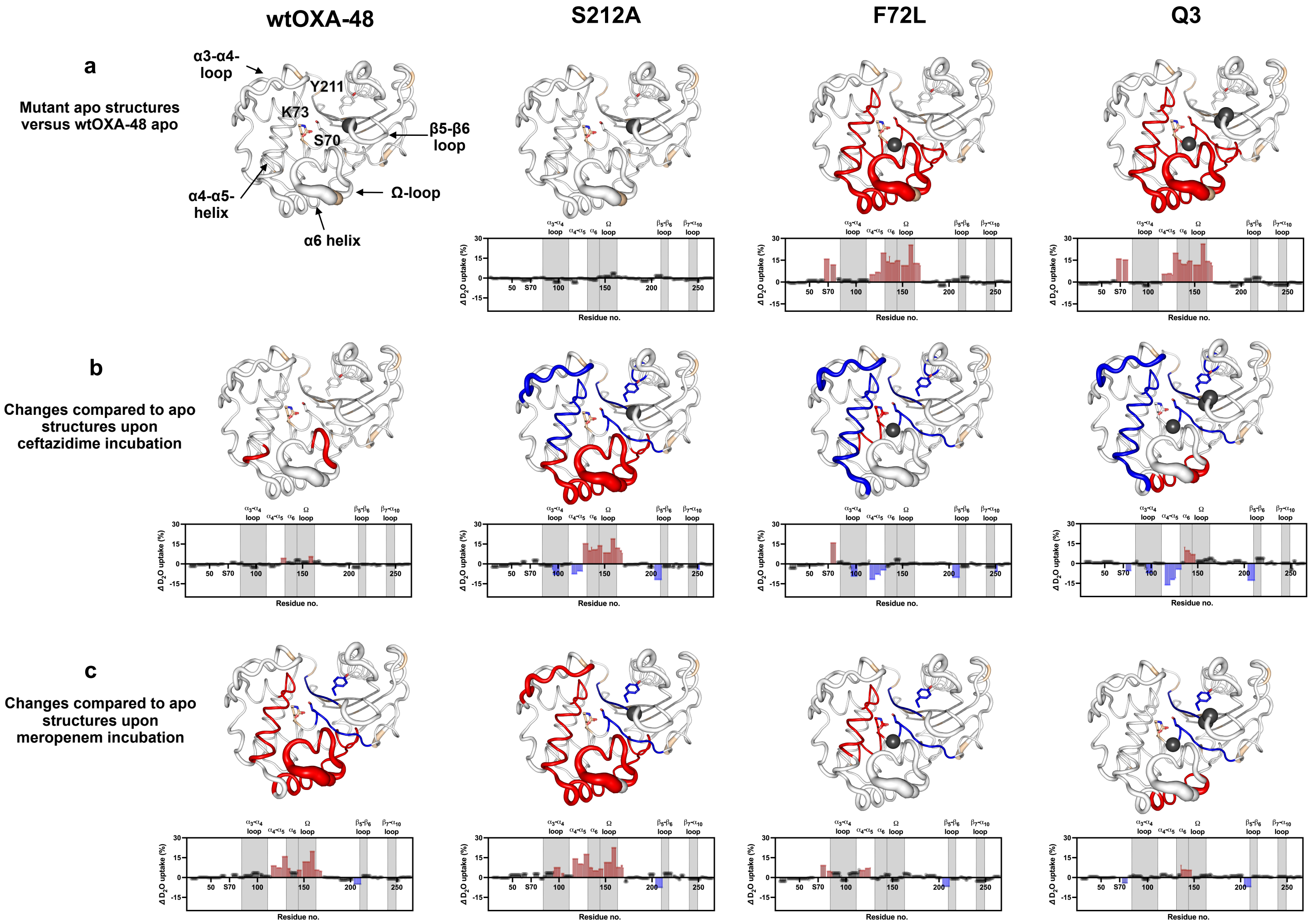


**Fig. S11: Effect of ceftazidime and meropenem turnover after 10 min HDX in the respective mutant backgrounds.**

Changes in $D_2O$ uptake after 10 min HDX for wtOXA-48, S212A, F72L, and Q3 using HDX-MS. For comparison, changes in $D_2O$ uptake induced by mutations are shown in panel a. Panel b (ceftazidime) and c (meropenem) describe changes during substrate turnover compared to changes inflicted by the mutants (panel a). Preincubation times were optimised based on kinetic parameters (Tab. 1 and Tab. 2) and results summarized in Tab. S3. For ceftazidime, the following substrate preincubation times were used: wtOXA-48 (60 min) , S212A (30 min), F72L (10 min), and Q3 (1 min). For meropenem, the following substrate preincubation times were used: wtOXA-48 (mix and inject) , S212A (mix and inject), F72L (1 min) and Q3 (1 min). For preincubation time optimization see Tab. S3. The relative increase (red) and decrease (blue) in $D_2O$ uptake (>|± 4|%) after 10 min HDX compared to the respective apo protein are mapped onto the structures. Black spheres display the location of the corresponding mutations (F72L, S212A, and T213A).

**a.** Panel a shows the effect of mutations on the $D_2O$ uptake compared to apo wtOXA-48. These variants were treated identically to the turnover experiments meaning that they were incubated for the same time frame but without substrate. No substantial changes were observed in the apo form of S212A compared to wtOXA-48. Apo forms of F72L and Q3 experienced a substantial increase in $D_2O$ uptake (15-30%) in the α4-α5 and α6 helices, the **Ω**-loop (peptide 148-156) and around the active site serine (peptides 72-78). Generally these results are in good agreement with our apo HDX experiments without preincubation time (Fig. S5)

**b.** Ceftazidime exposure resulted in limited changes in wtOXA-48, but induced up to 15% increased uptake within the **Ω**-loop and α6-helix of S212A. Decreased $D_2O$ uptake relative to apo forms in S212A, F72L and Q3 was observed in the α4-α5 and α6 helices and oxyanion hole peptide (with Y211), indicative of substrate binding in the active site. F72L and Q3 under ceftazidime turnover maintained enhanced $D_2O$ uptake compared to their apo forms in the α6-helix and **Ω**-loop, even showing further 10% increased uptake in the α6-helix of Q3. In addition, the region upstream and downstream of the active site serine S70 experienced an up to 15% increase in $D_2O$ exposure in F72L under ceftazidime turnover. While this region rigidified by ca. 4% in the ceftazidime exposed Q3 variants, it remained substantially higher than in the wtOXA-48 and S212A (Fig. 4c).

**c.** Meropenem exposure to wtOXA-48 and S212A led to enhanced $D_2O$ uptake in the α6-helix and **Ω**-loop, resembling the apo forms of F72L and Q3. Although changes in F72L and Q3 were less pronounced, a distinguishing feature of these variants was, similar to ceftazidime, the increased $D_2O$ uptake related to the region upstream and downstream of the active site serine S70 (peptides 72-78). This suggests that enhanced uptake within the active site region may relate to improved ceftazidime hydrolysis, while potentially compromising meropenem hydrolysis.

HDX= Hydrogen/deuterium exchange

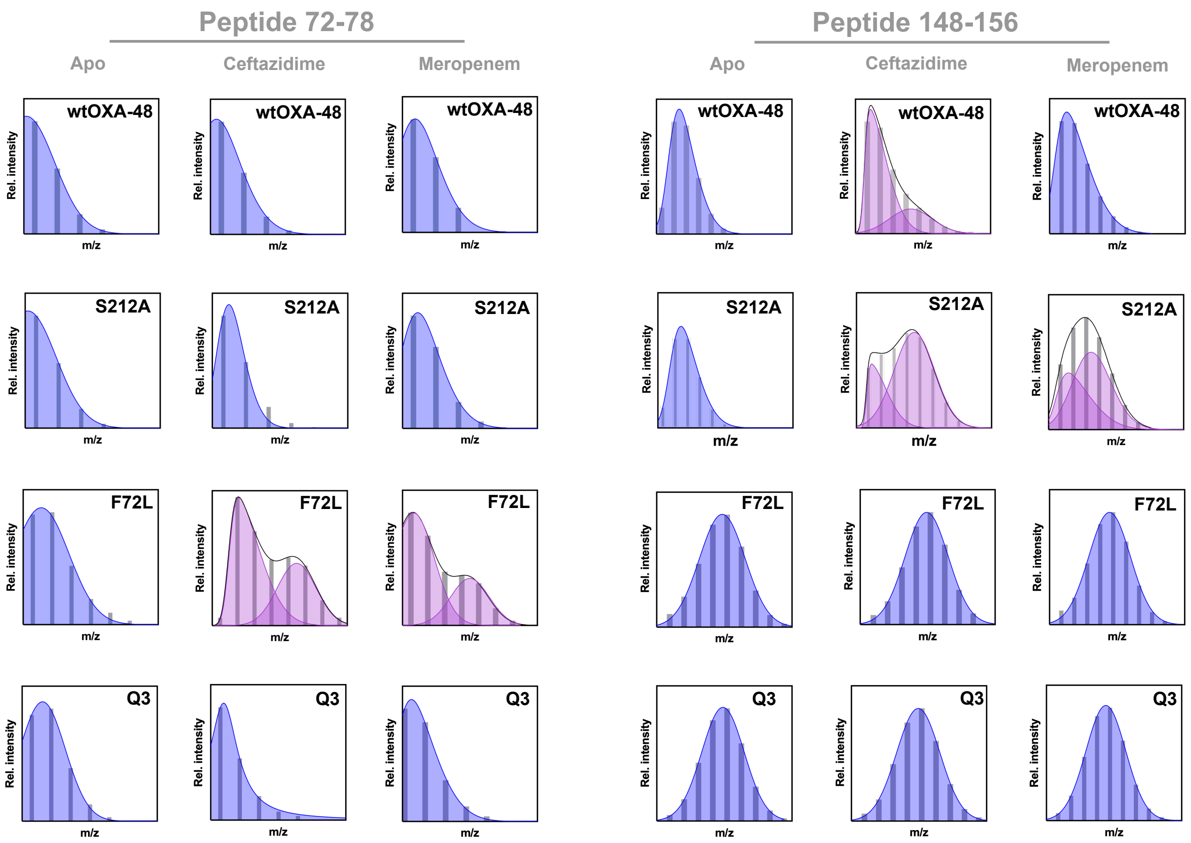


**Fig. S12: HDX-MS reveals the presence of two populations in F72L and S212A.**

Conformational distributions observed using HDX-MS after 10 min HDX for F72L (around active site, including the catalytic base K73: peptide 72-78) and 10 s HDX for S212A (partial Ω-loop: peptide 148-156) revealed conformational heterogeneity upon exposure to ceftazidime or meropenem. This heterogeneity likely reflects the evolutionary transition between distinct conformational states (Fig. 2 and Fig. 3). In addition, for the wtOXA-48 peptide 148-156 displayed ceftazidime-induced conformational heterogeneity. For ceftazidime, the following preincubation times were used: wtOXA-48 (60 min) , S212A (30 min), F72L (10 min), and Q3 (1 min). For meropenem, the following preincubation times were used: wtOXA-48 (mix and inject) , S212A (mix and inject), F72L (1 min) and Q3 (1 min). For preincubation time optimization see Tab. S3.

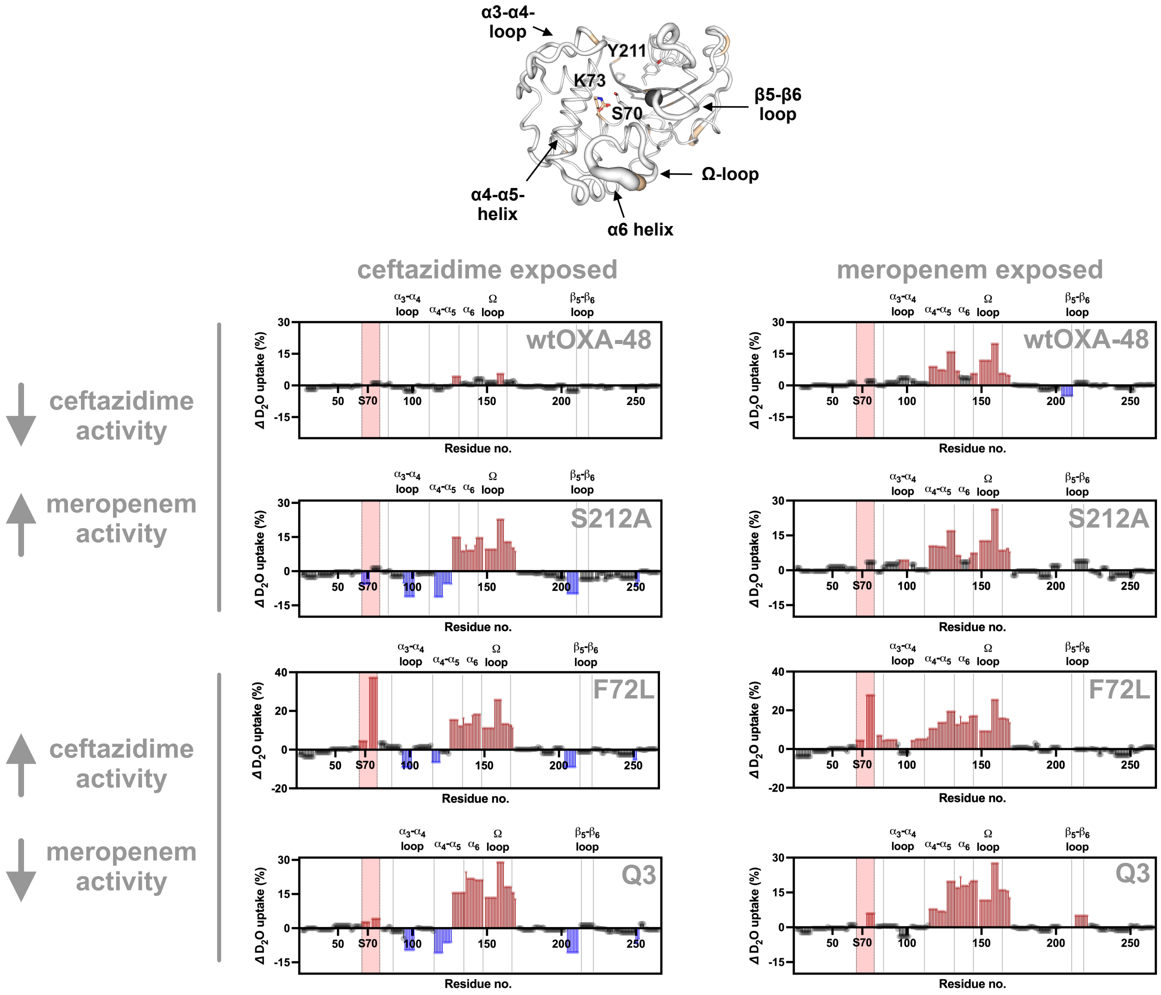


**Fig. S13: Overall $D_2O$ uptake between trade-off and non-trade-off variants.**

Changes in $D_2O$ uptake after 10 min HDX in substrate-exposed wtOXA-48, S212A, F72L, and Q3 compared to apo wtOXA-48. For ceftazidime, the following substrate preincubation times were used: wtOXA-48 (60 min), S212A (30 min), F72L (10 min), and Q3 (1 min). For meropenem, the following substrate preincubation times were used: wtOXA-48 (mix and inject), S212A (mix and inject), F72L (1 min) and Q3 (1 min). For preincubation time optimization see Tab. S3. Relative increase (red) and decrease (blue) are highlighted for regions > |± 4%|. Similar changes in $D_2O$ uptake relative to apo wtOXA-48 were observed in most of the structural regions, including the Ω-, β5-β6-and α3-α4-loops as well as the α4-α5-and α6-helices, for all mutants and both substrates. The region around S70 (peptide 72 to 78) displayed the highest increase in $D_2O$ uptake in the meropenem trade-off variants F72L (up to 35%) and remained elevated in Q3 (up to 5%). These effects were not observed in wtOXA-48 and S212A (for detailed statistics see Fig. 4c). All other regions behaved similarly, independent of the catalytic preference of the variants.

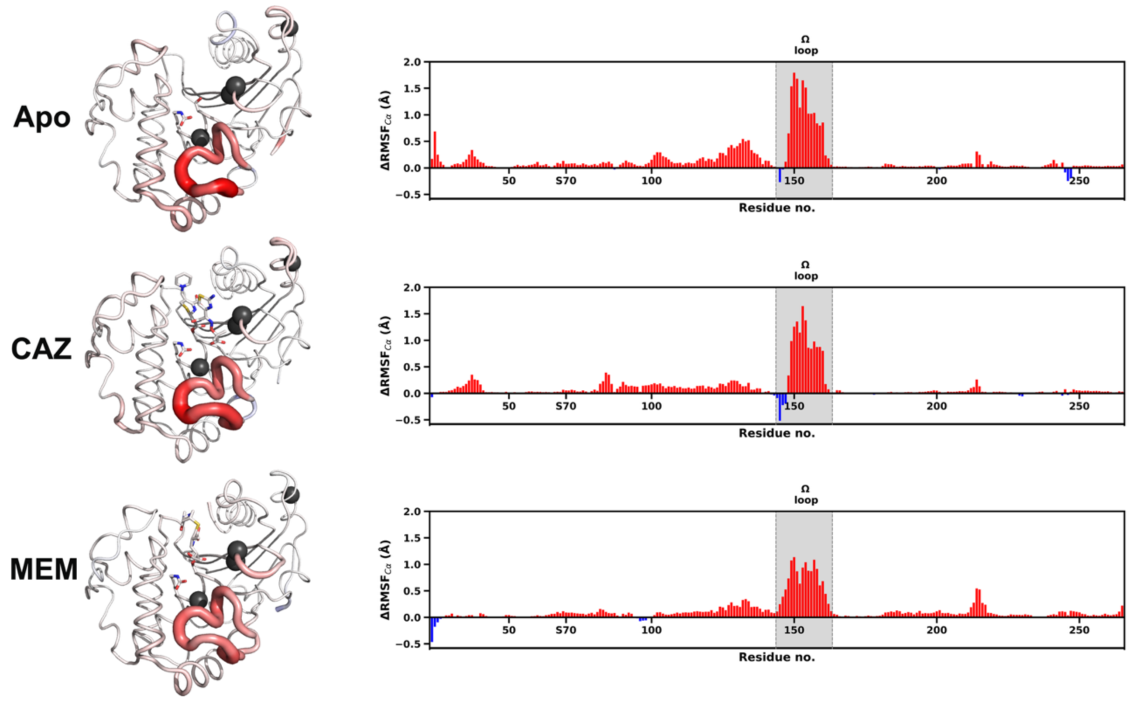


**Fig. S14: Comparison of the conformational flexibility between wtOXA-48 and Q4 in apo and acyl-enzyme complexes using MD simulations.**

⊿RMSF values based on Cα atoms of Q4 compared to wtOXA-48. Increase of $RMSF_{Cα}$ is coloured in red. The thickness and color of the tube are determined by ⊿RMSF values. Black spheres display the location of the corresponding mutations (F72L, S212A, T213A as present in Q3, and A33V, top right). A significant increase in the Ω-loop flexibility, similar to that observed in the HDX-MS data, was detected in Q4 in the apo, ceftazidime (CAZ)-bound and meropenem (MEM)-bound forms. Significance was determined by an independent two-sample, two-tailed t-test with Welch's correction, indicating $p<0.05$ for Ω-loop residues 148-162 in all three forms.

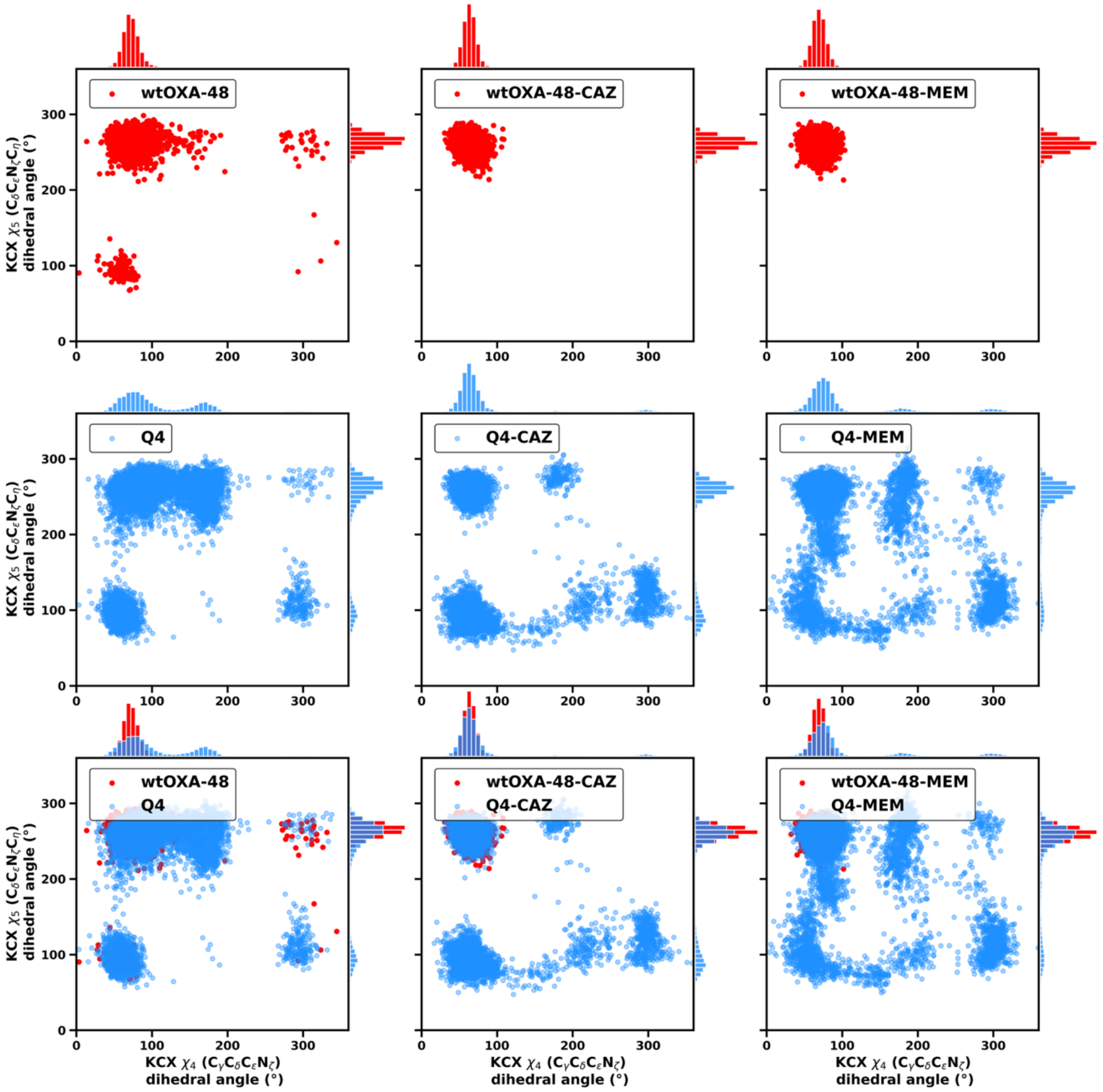


**Fig. S15: MD simulations display increased side-chain flexibility of the carbamylated K73**

The carbamylated K73 (KCX) side-chain $\chi$4 and $\chi$5 dihedral angles of wtOXA-48 (red) and Q4 (blue) sampled in the 30 ns simulations (n=32) of apo, ceftazidime-bound (wtOXA-48-CAZ and Q4-CAZ) and meropenem-bound (wtOXA-48-MEM and Q4-MEM) forms, shown as scatterplots with histograms (bin width: 5°).

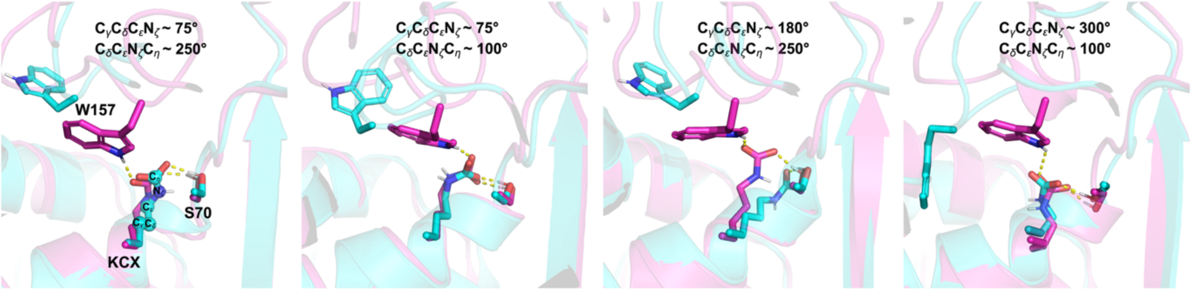


#### Fig. S16: Conformational freedom of the K73 side-chain

Representative snapshots of the different χ4 and χ5 KCX dihedral angles sampled in MD simulation, using wtOXA-48-CAZ (magenta) and Q4 apo (cyan) as examples. S70, KCX, the W157 side chains and ceftazidime are displayed as sticks and atoms used for dihedral angle measurements are shown as spheres. The wtOXA-48 hydrogen bond interactions between KCX and W157 are shown as yellow dashed lines. The frequency of the hydrogen bond interaction in the apo (0.0%), ceftazidime-bound (0.7%) and meropenem-bound (0.2%) is substantially lower than in the corresponding wtOXA-48 simulations (apo wtOXA-48: 56.0%, ceftazidime-wtOXA-48: 71.2% and meropenem-wtOXA-48: 57.6%).

### 3. Supplementary text

#### 3.1 HDX-MS: Extended method

##### 3.1.1 Sample preparation

**OXA-48 apo variants** were diluted in a 1:25 ratio (v/v) with labelling buffer (50 mM HBS, pD 7.2) to yield a final concentration of 1 μM enzyme in 95.9% $D_2O$. The reaction was then quenched after 10 s, 1 min, 10 min, 1 h, 4 h, and 16 h by addition of ice-cold quencher buffer (4 M guanidinium hydrochloride, 0.8% formic acid) in a 1:1 ratio, resulting in a final pH of 2.52. The last time point was performed at 37 °C for 3 h, corresponding to approximately 16 h of exchange at 20 °C.[6] This final time point was selected to explore slower dynamic processes without risking protein aggregation, which could otherwise interfere with accurate interpretation of the results.

**For substrate turnover experiments**, enzymes (10 μM) were incubated alone and with ceftazidime or meropenem (850 μM, see preincubation times in Tab. S3, Fig. S6 to S9). For these experiments (Fig. S6 to S9), a 10-fold dilution of both the apo and holo states was used to maintain a higher proportion of enzyme–substrate complex, yielding a $D_2O$ content of 90%. Preincubation time was optimized for wtOXA-48, F72L, S212A, and Q3 (Tab. S3, Fig. S6 to S9) based on the kinetic parameters for each variant (Tab. 1 and 2). Enzymes were incubated either with ceftazidime or meropenem, including no preincubation, or 1, 10, 30, and 60 min, before labelling in deuterated buffer for 10 s or 10 min. Substrate preincubation results that produced the strongest overall changes in conformational flexibility compared to their apo form were selected to compare changes in conformational flexibility, while prioritizing shorter preincubation whenever possible to minimize substrate hydrolysis (Tab. S3, Fig. S6 to S9). For instance, with the F72L variant incubated with meropenem (Fig. S9), a 1 min preincubation led to more pronounced changes in $D_2O$ uptake compared to 10 min, suggesting that extended incubation may cause excessive degradation of the substrate. In contrast, for wtOXA-48 with meropenem (Fig. S8), even without preincubation, substrate binding could already be detected, implying that longer preincubation may only increase hydrolysis.

##### 3.1.2 HDX-MS

Apo and holo quenched samples were subsequently injected into a Waters HDX Manager in-line with a Xevo G2S ESI Q-TOF (Waters, Milford, MO, USA) mass spectrometer. An Enzymate™ BEH Pepsin Column (Waters) was employed for on-line protein digestion at 15 °C and a flow rate of 100 μL/min. Generated peptides were then trapped on an Acquity UPLC BEH C18 VanGuard Pre-column (Waters) for 3 min, and eluted on an Acquity UPLC BEH C18 column (Waters) using a linear gradient of 2% to 35% acetonitrile (acetonitrile 100%, 0.1% formic acid) in 6 min. The flow rate was maintained constant at 100 μL/min. Several pepsin washes were performed after every injection to minimize carryover, using a 1.5 M guanidinium hydrochloride, 0.8% formic acid, 5% acetonitrile solution. To further minimize carryover, blank injections were also performed after each injection.

##### 3.1.3 Analysis

Peptic fragments generated from online digestion of unlabelled samples were identified with the aid of Protein Lynx Global Server 3.0 (Waters), while $D_2O$ uptake was analyzed with DynamX 3.0 software (Waters). Only peptides that were safely identified and commonly generated among all digested proteins were included in the experiment, with the following PLGS criteria employed to effectively choose high quality peptides for this study: (i) 1% retention time window in chromatographic separation, (ii) maximum error of 6 ppm for MH+ identification, (iii) 3 minimum fragment ions identified per peptide, (iv) 0.3 minimum products per amino acid, (v) 1 minimum consecutive product, (vi) only peptides whose length is between 4 and 30 amino acids. Fully deuterated samples used for back-exchange correction of peptides encompassing mutation sites were prepared.[7] Following pepsin digestion, approximately 65 overlapping peptides were consistently identified across all enzyme variants, resulting in complete sequence coverage (99%) and an average redundancy of 3.0.

Unless otherwise indicated, statistical significance of differences in $D_2O$ uptake at the peptide level was defined as a change $\geq |\pm 0.3|$ Da with a p-value threshold of 0.05, assessed using one-way ANOVA followed by a Tukey's post hoc test for all pairwise comparisons when more than two states were analyzed, or two-tailed t-tests when only two states were compared, as implemented in the DECA software package (github.com/komiveslab/DECA).[8] For residue-level analysis, obtained using DynamX heatmap scripts, a manual threshold was applied solely for visualization purposes, to visualize, on a continuous scale, the differences that were statistically significant at the peptide level. This approach was necessary because DynamX does not perform statistical validation at the residue level. Accordingly, for visualization purposes only, a 4% $D_2O$ uptake difference threshold was applied, ensuring that all peptides that had passed statistical validation were included while avoiding the inclusion of non-significant residues in the visualization.

#### 3.2 MD simulations: Extended method

For system building of the Q4-CAZ and Q4-MEM starting points (see main manuscript), Ω-loop residues E147-G161 of the Q5 apo form (PDB ID: 8PEB) were used, as preliminary simulations indicated consistency between MD ensembles and X-ray ensemble refinement. Q4-MEM additionally used the R214 conformation from this structure, to avoid instant formation of the R214-D159 salt-bridge (that then leads to Ω-loop ensemble not consistent with X-ray ensemble). For all models, crystal waters were retained where possible, and the carboxylate group of K73 (KCX) was added manually for both apo and holo forms of the enzymes in PyMOL v3.1.3. Protonation states of ionizable residues at pH 7.0 and histidine tautomers were determined by PropKa3.1 and the reduce program from AmberTools. To minimize the impact of protonation state differences on the results, one consistent set was used: all residues were in their standard protonation states, with all His residues being neutral and singly protonated on NE2. All complexes were solvated in a rectangular box of TIP3P water, with a minimum distance between the protein and the box edge of 10 Å. The solvated protein was neutralized with 1 or 2 $Na^+$ ions, depending on the system. Partial charges for meropenem acylated serine were obtained using the RED Server, based on HF/6-31G(d) RESP fitting.[9], with meropenem parameters obtained from on the General Amber Force Field 2 (GAFF2).

All systems were minimised in two steps. Both steps have 10000 steps of steepest descent followed by 10000 steps of conjugate gradient. In the first step, only the water molecules were allowed to move, all other atoms were restrained with a weight of 10 kcal mol $Å^{-2}$. Then, the temperature was increased from 50 to 300 K with pressure of 1 bar over a period of 20 ps. To maintain a similar conformation of the carbamylated lysine (KCX) for all systems before final simulations, the same restraints from previous work were applied only during the minimisation and heating procedure.[1] Flat-bottom restraints for side-chain KCX dihedral angles ($C_\delta C_\varepsilon N_\zeta C_\eta$ (≤ 230° and ≥ 260°) and $C_\gamma C_\delta C_\varepsilon N_\zeta$ (≤45° and ≥ 95°) were applied for all systems. For acyl-enzyme systems, the distance restraints from the deacylating water to either the KCX base oxygen or the antibiotic electrophilic carbon were applied too. All restraint force constants were 10 kcal mol $Å^{-2}$ during the heating and 100 kcal mol $Å^{-2}$ during the minimisation. During all equilibration and production simulations, periodic boundary conditions were applied, and the SHAKE algorithm was applied to fix all bond lengths involving hydrogen atoms. A time step of 2 fs was used and cutoff radius for non-bonded interactions was set to the default 8.0 Å. For all systems, 32 independent simulations of 10 ns of equilibration followed by 30 ns of production in the isothermal–isobaric ensemble

(NPT) ensemble at 300 K and 1 bar were carried out (temperature controlled with Langevin dynamics, collision frequency 0.2 $ps^{-1}$, pressure with Berendsen barostat, pressure relaxation time 1 ps). No restraints were applied during the equilibration and production stages. All simulations were conducted using the Amber24 package (Case et al., 2024).

The production trajectories of simulations were analysed using CPPTRAJ (Roe et al., 2013). Root mean square fluctuations (RMSF) values based on Cα atoms ($\Delta RMSF_{C\alpha}$) were calculated by using RMSF per residue of each OXA-48 variant subtracted by the RMSF value of corresponding residue in wtOXA-48 (Tab. S4). RMSF values of each residue were determined by first getting the average coordinates of each replica and then align trajectory to these coordinates to compute RMSF values. The KCX dihedral angles ($C_{\delta}C_{\varepsilon}N_{\zeta}C_{\eta}$ and $C_{\gamma}C_{\delta}C_{\varepsilon}N_{\zeta}$) were measured, and hydrogen bond analysis was performed with a bond angle cutoff of 135° and a distance cutoff of 3.0 Å between donor and acceptor. Structure images were made in PyMOL.

428
